# Evaluating microbial contaminations of alternative heating oils

**DOI:** 10.1101/2022.09.04.506525

**Authors:** Maximilian J. Surger, Katharina Mayer, Karthik Shivaram, Felix Stibany, Wilfried Plum, Andreas Schäffer, Simon Eiden, Lars M. Blank

**Author notes:** Corresponding authors Prof. Dr. Lars M. Blank and Dr. Maximilian J. Surger, Institute of Applied Microbiology RWTH Aachen University Worringer Weg 1, D-52056 Aachen.

## Abstract

Since 2008, legislative initiatives for climate protection and reduced dependency on fossil resource imports led to the introduction of biofuels as CO_2_-reduced alternatives in the heating oil sector. In the case of biodiesel, the oil industry or its customers were confronted with accelerated and escalating microbial contaminations during heating oil storage. Since then, other fuel alternatives, like hydrogenated vegetable oils, gas-to-liquid products (GtL), or Oxymethylenether (OME) have been or will be developed and potentially introduced to the market. In this study, we use online monitoring of microbial CO_2_ production and the simulation of onset of microbial contamination to investigate the contamination potential of fuel alternatives during storage. As reference and blends, fossil heating oils of various refineries, in the course of this from various crude oils, and refinery processes reveal considerable variation in potential microbial activity. Oxymethylene ethers have an antimicrobial effect, while various forms of biodiesel confirm the promotion of microbial activity and diversity. The paraffinic Fischer-Tropsch products and biogenic hydrogenation products demonstrate high resistance to microbial contamination despite allowing microbial diversity. Through an array of analytics, including advanced chromatography coupled mass spectrometry, elemental analysis, and microbial sequencing, we can discuss critical fuel properties that promote or inhibit microbial contaminations. In summary, novel, non-fossil heating oils show different strengths and weaknesses for long-term storage. Designing blends for microbial activity reduced long-term storage might be an option. While being niche products, these fuels will contribute to the rapid reduction of fossil resource use.

## 1. Introduction

### Microbial contamination during heating oil storage

Ten million households and twenty million people in Germany, approx. 25% of the population, are stably supplied with heat via heating oil EL (extra light). The industry expects that in the course of the Climate Protection Plan 2050, fossil fuels will be successively replaced by more climate-friendly alternatives, but that the importance of liquid fuels will be maintained due to the existing infrastructure, their energy density, and storage stability [1–5]. However, this storage stability is increasingly discussed critically by consumers and industry. In addition to abiotic aging, the focus is primarily on the susceptibility to microbial contamination and accompanying damage (i.e., biofouling) of burner systems [6]. This is not a new phenomenon, for example, even before the introduction of FAME the involvement of microorganisms was documented for 34% of the corrosion events in the mineral oil industry before 1999 [7].

The consumption of mineral oil (predominantly heating oil EL) to supply heat to households and businesses fell to below 24 million t in Germany by 2018 [2–4]. The decrease in consumption of heating oil EL correlates with an increase in storage time due to the technical development of burner systems and improved thermal insulation. Currently, heating oils are stored by the end customer for at least half a year up to a maximum of five years, which depends on seasonal purchase and storage capacity. The application of modern condensing systems increases the storage time by another 30%, and the application of hybrid systems, i.e. the combination with wood log boilers or solar collectors with water storage, increases the storage period by 100% [5, 8]. The decrease in consumption and advancing application of hybrid systems are a clear indicator of the transition from tank and burner systems to substitute infrastructure or emergency power systems. So that the tank or burner systems established in millions of households will continue to be used for decades, but of course increasingly with alternative non-fossil fuels. Due to a continuous increase in storage time in private households occurrence and intensity of microbial contamination of heating oil EL will increase and the issue will only continue to grow in importance for the industry [9]. Increasing interest is also being shown in the topic for national oil and gas reserves. The EU states, but also the USA, Japan, and Australia, maintain huge oil and gas depots to meet their obligation to supply the population for 90 days in exceptional situations.

Fundamental to the outbreak of microbial contamination in heating oil storage is the presence of water, mainly by the penetration of atmospheric moisture into the tank via the tank ventilation that is required for technical reasons. Exceeding the water absorption capacity of the fuel (for fresh fossil heating oil EL approx. 100 mg/kg) free water phases are formed, which may be stably present in the oil forming emulsions, but may also settle or break depending on temperature fluctuations (seasonality of contamination events) and surface tension, resulting in the formation of a separate free water phase at the bottom of the tank. Emulsions or separate water phases with volumes of 1-3 µl and water activity (a_w_) of >0.8 are the basis of microbial contamination [10–19].

Further progression of microbial contamination is significantly influenced by the availability of nitrogen. Field reports document 13 to 1,300 mg/kg of nitrogen in heating oil [11–12, 19] and 95 mg/L (101 mg/kg at a density of 0.94 g/cm^3^) in crude oil [15] in the form of ammonium salts and heterocycles such as anilines, pyridines, quinolines (alkaline) or indoles, and carbazoles (non-alkaline) [20]. The main reason for the variation in reported nitrogen content is the use of desulfurized or non-desulfurized fuel. When the fuel is treated with hydrogen at high pressure, the bound nitrogen is removed at the same time [21]. A limitation of microbial diversity and the enrichment of nitrogen-fixing genera such as *Pseudomonas* is an indication of a shortage of nitrogen components [22].

Other factors that drive microbial contamination are osmotic pressure, salinity, oxygen availability, carbon sources (hydrocarbons), temperature, and pH [19].

### In field assessment of microbial contamination in mineral oil products

If possible, a test to fully assess microbial risk must cover all three parameters of microbial viability, i.e., biomass development (growth), metabolic activity, and membrane integrity [22, 23–24]. The use of classical methods, such as measurement of optical density, dry cell weight (CDW), and determination of colony-forming units (CFU) in heating oil storage are challenging, as representative sampling in such an oil/water phase setup is non-trivial. Classic methods consider mainly the biomass formed, ignore cultivability in the laboratory, are affected by emulsions and oil droplet adhesion, and rely exclusively on single invasive sampling in predefined tank areas. Nevertheless, the determination of colony-forming units (CFU) is significantly used today to decide whether contaminated heating oil can still be used, based on numerous standards (IP 385, IP 613, ASTM D7978-14, ASTM D6974-20, and DIN 51441) [16, 25–30]. Increasingly important for the assessment of microbial contamination in practice is the adenosine triphosphate (ATP) assay, for which some application standards have already been formulated (ASTM D7463 - 21 and ASTM D7687 - 21). The ATP assay takes into account biomass, metabolic activity, and, when extracted from whole cells, membrane integrity. Further, classical cultivation in the laboratory is not required. Available assay kits offer maximum sensitivity (up to 1 pmol/ml ATP) and cross-species applicability. Remaining disadvantages are the single invasive sampling from predefined tank areas, the adherence of oil, and the overall susceptibility of the assay system to the sample matrix [16, 31–36]. Various systems for permanent online monitoring of the microbial status in heating oil tanks, such as tracking of oxygen consumption or detection of chemical gradients considering biomass and metabolic activity parameters, are under scientific discussion [7][34]. In previous work, non-invasive online monitoring of microbial CO_2_ production during the simulation of a contaminated heating oil tank was introduced as a tool for seamless tracking of microbial contamination and will be a key tool in the here presented study [37]. Microbial CO_2_ production as a measure of biomass and catabolic activity has been successfully used by Zhang et al. (1998) to measure the degradability of diesel or by Rose et al. (2020) to measure the degradability of plastic monomers based on single invasive sampling [25, 38–39].

### Current organizational, technical, and chemical methods for microbial control in oil tanks

The current and relevant standards for heating oil EL (DIN EN 51603-1 and DIN SPEC 51603-6) describe the physico-chemical requirements for the fuel. Neither relevant requirement parameters for microbial activity are defined in them, nor are limit values specified (for example a maximum nitrogen content or acid number). So far mineral oil companies in Germany are not legally liable for microbial contamination as a result of water entering the heating oil tank and there is currently no willingness of the politics or technical committees to impose legal requirements (for example a maximum nitrogen content or acid number) on the microbial stability of heating oils [40–41]. The technical standard on substances hazardous to water (TRwS) 791 for heating oil consumer installations and the directive on installations for handling substances hazardous to water (AwSV) concretize the requirements of the Federal Water Act (§62 para. 4 WHG) and the water laws of the states (WG). Of particular relevance for microbial stability are the obligations of private operators formulated therein to control infiltrated water (including leak protection, fill levels, and bulges). Maintenance intervals of 2.5-5 years by expert maintenance companies and defect reports by supplier companies following refilling are recommended. These time dimensions already allow the macroscopic manifestation of microbial contamination in the form of biofilms and corrosion [14, 42–44]. However, the reality is the lack of widespread control, reliance on responsible maintenance, and reactive maintenance. This is when voluntary recommendations of the maintenance industry for microbial limits based on colony-forming units (CFU/L) from defined tank areas and recommendations for removal of infiltrated water, temporary pump-down (heating and mixing), filtration of fuel oil (≤ 1 mm pores with no claim of sterile filtration), mechanical removal of deposits, filter replacement, and varying intensity of biocide use apply. Not uncommonly, the operator experiences the renewed biofilm-associated clogging of filters a few months later [9, 12, 16, 45].

The focus of technical innovations in heating oil tanks or burner systems with relevance to microbial activity is also on controlling the water balance and separating the water and oil phases. Thus, the change from two-line to single-line suction avoids backflow of thermally stressed heating oil and in this course microbially easier accessible carbon sources into the tank and corresponding turbulence of the tank bottom with potential water phase and microbes. In addition, there can be no entry of metal ions or trace elements from corrosive lines. A deflected outlet in the two-line system reduces turbulence. The introduction of floating suction, which always rests on the surface of the oil phase, ensures the maximum distance to the potential water phase with biofilm resting on it [45–46]. Drying systems, such as water separators, are used to sustainably control the water content in the oil phase itself and can keep the water concentration at < 60 ppm [12].

The focus of chemical measures for microbial control is certainly on the use of biocides (antibiotics or fungicides) as additives in liquid fuels or as tank coatings (e.g., rifampicin, minocyclines, cefazolin, sulfadiazines) [47]. The efficacy of biocides depends on multiple factors such as half-life, concentration, contact time, the extent of existing contamination, presence of biofilm/sludge/sediment, pH of the water phase, presence of inactivating substances such as sulphides, temperature, partition coefficient between water and oil phases, and water to oil phase ratio [16, 47]. The intensity of use in practice ranges from the highest doses (shock biocide dosing) to regular low doses, directed by the extent of the infestation, available time, and risk of resurgence [16]. However, the widespread and prophylactic use of biocides is also critically discussed in the storage of liquid fuels. In addition to the risk of selection of resistant strains and the persistence of still problematic dead biomass, many biocides have lost their approval or the approval of new biocides is more difficult with the enactment of the REACH regulation in 2007 [48]. Biocide-independent strategies partly build on existing additives that have indirect antimicrobial activity. Antifreeze agents distribute mainly in free water phases, provide high dissolved concentrations, and a strong reduction of water activity (a_w_) [16]. Surfactants are a double-edged option. Lowering surface tension and suppressing coagulation of cells certainly suppresses biofilm formation as the primary annoyance of microbial contamination. However, the overall microbial activity of single cells (including the secretion of corrosive acids) is not reduced, and the spread of single microbes in the tank and burner system is even promoted [17, 47]. Growth-promoting compounds can be specifically withdrawn from the access of microbes. Installing a reservoir with desiccant (e.g., calcium chloride) can reduce the entry of atmospheric moisture. The application of Lactoferrin that binds iron, which is essential for bacterial growth, has to be investigated [16, 47].

### Use of alternative fuels

The EU Directive (REDII, 2018/2001) on the promotion of the use of energy from renewable sources and the “Renewable Energies Heat Act” (EEWärmeG, October 2015) demand a renewable share of 14% of liquid fuels and an increase in the renewable energy share for heat by 1.1% per year by 2030 [49–50]. In 2019, the German government even formulated a 66% reduction of CO_2_ emissions in heat supply for buildings in the “Climate Protection Plan 2050”. In addition, liquid fuels were subject to additional costs in the form of CO2 certificates from 2021 under the Fuel Emissions Trading Act (BEHG) [4]. To continue to use the existing predominantly private infrastructure for liquid fuels and to permanently store energy generated from solar, wind, and hydropower (“power-to-liquid (PtL)”), liquid fuels continue to exist [8, 51]. In this context, fossil fuels are being potentially replaced by new alternative CO_2_-reduced or -neutral biogenic or synthetic fuels, which can be distinguished based on feedstock, manufacturing pathway, and possibly microbial susceptibility. As a result, there is currently an enormous diversification of the fuel market before alternative standard blends will become established in individual application areas [4]. For the application of biodiesel in the heating oil sector, the DIN EN 14214, DIN SPEC 51603-6, and DIN 51603-1 (FAME- free) standards were last updated in 2017-2019, which define EL heating oil with proportions of 0-100% and specify requirements for operational safety in certain burner systems [40–41, 52]. A technical specification (DIN/TS 51603-8) is currently (2021) being worked on for the application of paraffinic fuel oil alternatives [53]. The environment of heating oil storage, in which microbial contaminations develop is thus subject to massive changes concerning substrate availability or diversity, water solubility, and toxicity.

### Relevance of Biodiesel for Microbiology

Biodiesel or fatty acid methyl esters (FAME) are produced via transesterification of vegetable oil-based triglycerides and methanol. By-products are phospholipids and free fatty acids. Antioxidants (in ppm) are added and plant fertilizers are present in traces. The energy content of 1 L of biodiesel corresponds to the energy content of 0.9 L of diesel [54]. In Germany, the most important feedstock in 2021 was recycled used cooking oil (37%, Used Cooking Oil, UCO), followed by rapeseed oil (33%), palm oil (25%), and soybean oil and sunflower oil (1% and 3%, respectively) [55]. Celebrated for its environmental friendliness due to generally low emissions, especially CO_2_ neutrality and biodegradability, FAME soon revealed limited long-term storage stability in heating oil applications due to low autooxidation resistance and biodegradability or microbial susceptibility, resulting in biofouling [56–57]. Increased biomass formation, increased degradation rates, and an increase in microbial diversity in the form of greater proportions of yeasts and molds have been widely documented with biodiesel blends [12, 14, 16, 18, 33; 57-59]. In the research report 611 of the DGMK from 2002, the author concluded that the addition of only 5% biodiesel, should result in shortening storage of heating oil to six months [60]. Currently, the following standards DIN EN 14214, DIN SPEC 51603-6, and DIN 51603-1 exist, which define heating oil EL Bio with 0-100% biological fuel components and allow it for certain burner systems [40–41, 52]. In Germany, a realistic future market target is the blending of up to 20% biodiesel to heating oil EL.

One reason for microbial susceptibility is the influence on the water balance in fuel oil storage. Depending on the degree of saturation of the fatty acid methyl esters, biodiesel has a strong hygroscopic behavior [12–13][61] and due to the polar molecule sections, the overall water absorption capacity is also increased. A high water content, linearly dependent on the biodiesel content (Table 1) [16, 59], ensures the survival of invading microorganisms [13]. As soon as microbes could settle, their metabolism of highly reduced heating oil molecules results in additional water formation. However, emulsions and separate free water phases are formed only after the water absorption capacity has been exceeded, which causes microbial infestation to progress. The addition of biodiesel therefore initially delays this progression.

**Table 1:**
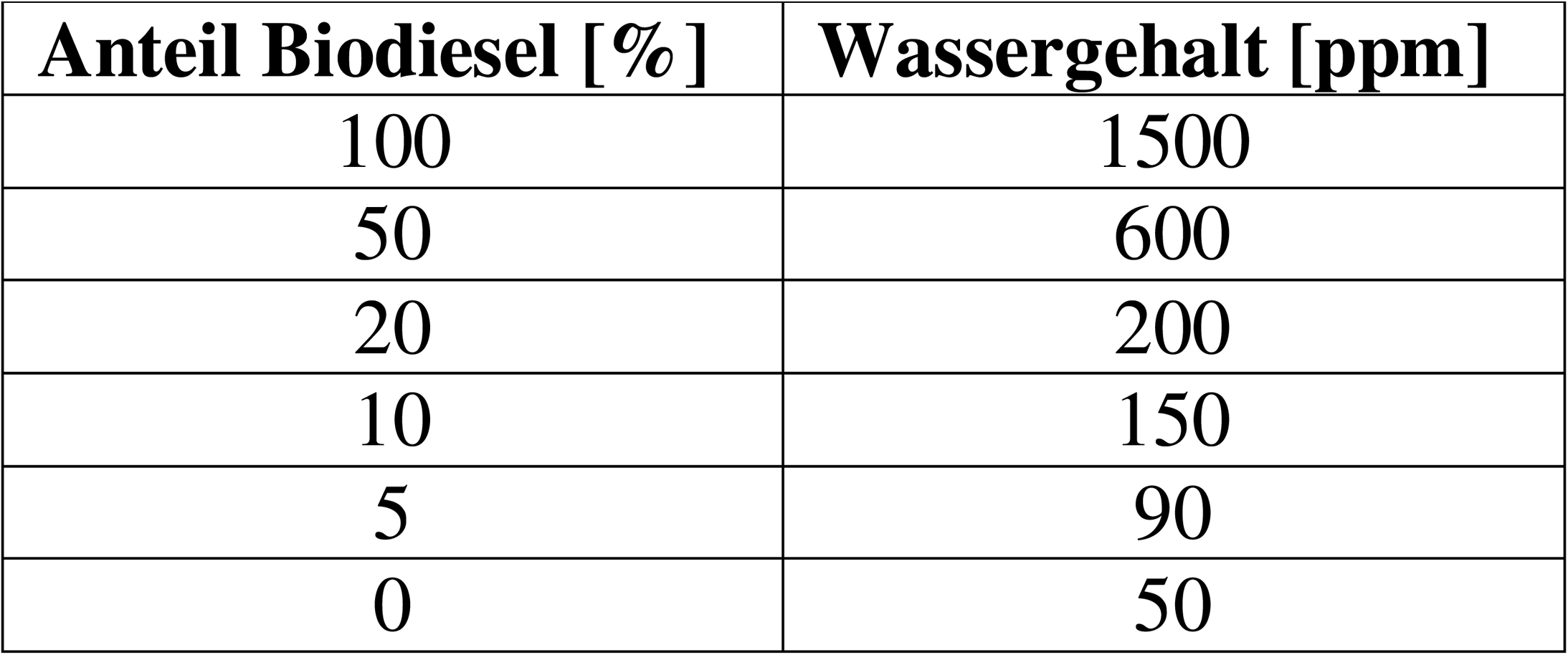
Water content of heating oil depending on share of biodiesel. [60].

As surface-active substances, the fatty acid methyl esters lower the surface tension of the water, leading to increased emulsion formation. The stabilized emulsion allows microorganisms to contaminate the fuel phase itself, independent of a free separated water phase, and in the case of heating oil, to spread more widely in the storage tank and burner system [9, 11–13, 16–17, 59]. Another reason for the increased microbial susceptibility is the improved elemental composition or decarbonization of the fuel. Pure EL (low sulfur) heating oil consists of 85.5% carbon, 13.5% hydrogen, and 0.56% oxygen, while pure biodiesel consists of 55.6% carbon, 19.4% hydrogen, 11.7% oxygen, and 2.7% nitrogen. The increase in oxygen content is due to the molecular class of fatty acid methyl esters and their starting and decomposition products. The input of nitrogen and phosphate is due to nitrogen- containing antioxidants, e.g., phenylene diamine, and phospholipids as a byproduct of biodiesel production, as well as fertilizer contamination [11][13][16][62]. An optimized ratio of carbon to nitrogen to phosphorus (C:N:P) promotes the degradation of each carbon source and the build-up of new microbial biomass with an average chemical composition of C (48%, w/w) O (24%, w/w) N (14%, w/w) H (7%, w/w) P (3%, w/w) S (0.5%, w/w) [63–64]. Carboxylic acids of varying chain length and short-chain alcohols occur as by-products of biodiesel production, as secondary microbial metabolites, and as additional simple carbon sources, as documented by elevated acid numbers [52, 65]. Apart from non-targeted nutrients, FAMEs themselves represent simple and energy-rich carbon sources for microorganisms depending on chain length and degree of saturation (soybean/rapeseed: C18-22, unsaturated > coconut/palm: C12-14, saturated). The degradation of FAMEs requires only hydrolytic enzyme activities (like lipase and esterases), which are common microbial tools, whereas the degradation of hydrocarbons requires specialized oxygenases and conditions that allow for high oxygen consumption [6, 9, 16, 34]. As a consequence of the addition of fatty acid methyl esters as a simple microbial nutrient source, synergistic degradation of the fossil fraction of the fuel mixture has also been widely documented and described (co-metabolism). Possible reasons for the improved utilization of generally inert heating oil are numerous and here only briefly mentioned. The FAME or other second substrates increase the rate of degradation of the first substrate, because the second substrate increases population size, or degradation of the second substrate produces stimulatory growth factors such as biosurfactants, or the degradation pathways share individual enzymes, or the degradation of the second substrate provides energy for the expression of the degradation pathway of the first substrate [14, 56–57, 66]. Other reasons for microbial susceptibility include reduced oxygen diffusion (1.55%/h for pure fuel oil compared to 1.33%/h for fuel oil with 20% (v/v) rapeseed oil methyl ester (RME)) and longer contact time of oxygen with the fuel. In addition, a higher content of acidic and reactive groups promotes abiotic aging of the fuel and corrosion events on the tank, so that additional nutrients are released over time, and niche and sediment formation occurs. This is regularly documented by fuel storage stability indices, i.e., increased acid numbers and decreased oxidation stability [8-9, 11, 17-18, 52, 59, 67-68].

### Microbial relevance of paraffinic fuel alternatives

The paraffinic fuel alternatives currently include hydrogenated vegetable oils (HVO, biogenic) and the products of Fischer-Tropsch synthesis (XtL, synthetic) [8]. The production of hydrogenated vegetable oils can be derived from the same feedstocks as the production of biodiesel (FAME) [69]. However, due to the manufacturing process, the product properties are less dependent on the properties of the raw materials, so manufacturers, following economic and environmental aspects, increasingly rely on palm oil (high saturation level requires less hydrogen in the manufacturing process, world market availability) and used cooking oils (CO_2_ balance) as raw materials [70–75]. HVOs are produced in two steps.

Hydrogenation saturates double bonds with hydrogen and removes oxygen from ester bonds by splitting off water or CO_2_ [70]. Unbranched alkanes with 15 to 18 carbon atoms are formed, corresponding to plant fatty acids. In a second step, most of the alkanes are converted by isomerization into iso-alkanes with advantageous physical properties. The energy content of 1 L of HVO replaces 0.96 L of diesel [8, 70, 76–81]. The starting material of the Fischer-Tropsch process is synthesis gas consisting of hydrogen and carbon monoxide. Depending on the origin of the synthesis gas, it is referred to as BtL (biomass to liquid), PtL (power to liquid), GtL (gas to liquid), and CtL (coal to liquid). The products of the Fischer-Tropsch synthesis are very long-chain unbranched aliphatics. However, with increasing chain length, branched and to a lesser extent, cyclic hydrocarbons are already produced [82]. Hydrocracking and hydro-isomerization also yield predominantly branched alkanes with chain lengths in the middle distillate range. The energy content of 1 L of BtL is equivalent to 0.94 L of diesel [8, 69, 81, 83–86]. The reality for the use of paraffinic fuel alternatives in the heating oil sector (heating oil EL P) is a draft technical specification DIN/TS 51603-8, which in principle allows replacing fossil heating oil EL to 100% by paraffins. The technical limitation is the reduced lubricity and density due to the lack of aromatics. This could be compensated by blending with biodiesel, which is currently not permitted. A market-realistic target is the blending of 50% paraffinic fuel alternatives to fossil EL heating oil or 20% biodiesel and 50% paraffinic fuel alternatives to fossil EL heating oil [53, 81, 87–88]. The paraffinic fuel alternatives are celebrated for high cetane numbers, high ignition readiness, clean combustion, and thus generally low emissions [88–90]. From a microbial perspective, the addition further lowers the water affinity and content of fossil heating oil, which limits microbial survival in the oil itself. The surface tension of the free water phases remains unaffected, so no stabilization of emulsions and reduced spread of existing microbial contamination in the storage tank or burner system is expected [81]. Nevertheless, separate free water phases are expected to form even faster via condensation. The DIN/TS 51603-8 provisional technical specification also indicates a nitrogen content limit of 140 mg/kg, implying high nitrogen availability, optimized carbon to nitrogen ratio, and more efficient degradation of the carbon source [53, 81]. The admixture of paraffinic fuel alternatives increasingly limits the diversity of microbial carbon sources to a single challenging carbon source. Also relevant is the reported high abiotic storage stability, so less corrosion, niche, and sediment formation is expected compared to biodiesel [73, 81]. Practical empirical data and scientific studies on biodegradation performance, biomass development, microbial diversity, and effects of contamination are currently not documented.

### Microbial Relevance of Oxymethylenethers

The starting product of oxymethylene ether (OME) synthesis is methanol, which is produced from synthesis gas. OME is produced via formaldehyde or trioxane via polymerization reactions with further methanol and formaldehyde. The oxymethylene ethers with the chemical structure H_3_C-O-(CH_2_O)_n_-CH_3_ (OME_n_) contain no C-C bonds, carbon atoms are connected via ether bridges. For application as an alternative fuel in the diesel/heating oil sector, a blend of the oligomers OME_3-5_ is used and an admixture of up to 15% (v/v) is targeted [91–96]. The high oxygen content ensures cleaner combustion and lower emission levels even when small proportions are blended with fossil fuels, but it is also responsible for a low energy content [92, 94–98]. The energy content of 1 L of OME replaces approx. 0.5 L of diesel. Good microbiological degradability of related oxygenates such as methyl tert-butyl ethers (MTBE), for example, by *Pseudomonas* sp. is reported and their environmental compatibility, which can certainly be applied to OMEs [99–101]. These reports are countered by the problem of oxidative decomposition reactions and acid-catalyzed back reactions of OMEs to the toxic or carcinogenic starting products methanol and formaldehyde [102]. The oxidative decomposition reactions are radical chain reactions involving oxygen radicals triggered by light and temperature. The acid-catalyzed reverse reaction is determined by the availability of the co-product water of OME production [95, 98]. Practical experience and scientific studies on biomass development, microbial diversity, and effects of contamination are not documented at present.

### Objective of this study

We investigated the influence of different refineries and blending of the latest alternative fuels on microbial susceptibility in heating oil storage. For that, we used continuous online monitoring of microbial CO_2_ production as a measure of microbial activity, a microbial mixed culture representative for the heating oil sector, and a simulated onset of microbial contamination. While FAME used in heating oil storage fostered contaminating microbes, formation of biofilms, and supported corrosion events, the newer fuel alternatives are microbially less vulnerable. Crucial properties of the fossil heating oils or alternative fuels for promoting or inhibiting the microbial contamination process were revealed. In addition to online monitoring of microbial activity, the monitoring of other indices of oil and infiltrated water were identified as tools for early detection of an advancing contamination process.

## 2. Materials and Methods

### 2.1 Microbial strains and storage cultures

To mimic an oil-storage-tank (1:5 approaches) 50 ml (100 ml in case of OME blend investigation) of the free water phase, consisting of 0.1% NaCl, were overlaid with about 250 ml (500 ml in case of OME blend investigation) fossil heating oil or alternative fuel within a 500 ml shot bottle (1L shot bottle in case of OME blend investigation), resulting in a 300 ml (550 ml in case of OME blend investigation) headspace. To investigate the effect of microbes on fuels 50 ml of free water phase were overlaid with 10 ml fossil heating oil or alternative fuel within a 100 ml glass bottle with constant air supply via needle and sterile filter (5:1 approaches) (Figure 1). A volume of 800 ml water phase was inoculated with a mixture of 20 representative heating oil microbes as defined by Leuchtle et al. [25] including an amount per strain corresponding to 16 mg CDW (a total of 320 mg CDW for all microbes). The microbial mixture did not contain anaerobes as oil storage tanks are ventilated [25]. The microbes used are found in Table 2. The precultures were made in LB (10 g/L peptone, 10 g/L yeast extract, 5 g/L NaCl medium), YEP medium (20 g/L peptone, 20 g/L glucose, 10 g/L yeast extract), YEPS medium (10 g/L peptone, 10 g/L yeast extract, 10 g/L sucrose), potato extract glucose bouillon (PEGB, 26,5 g/L), or ME medium (20 g/L malt extract, 1 g/L peptone) for bacteria, yeasts, *Ustilago maydis*, *Rhodotorula mucilaginosa*, and molds, respectively. Precultures of single strains consisted of 100 ml medium in 1 L Erlenmeyer flasks without baffles grown into early stationary phase. Those single cultures were shaken at 200 rpm and incubated at 30°C. The precultures were washed with 0,1% NaCl before use. The shot bottles were not shaken, were kept in the dark, and at room temperature.

**Figure 1.**
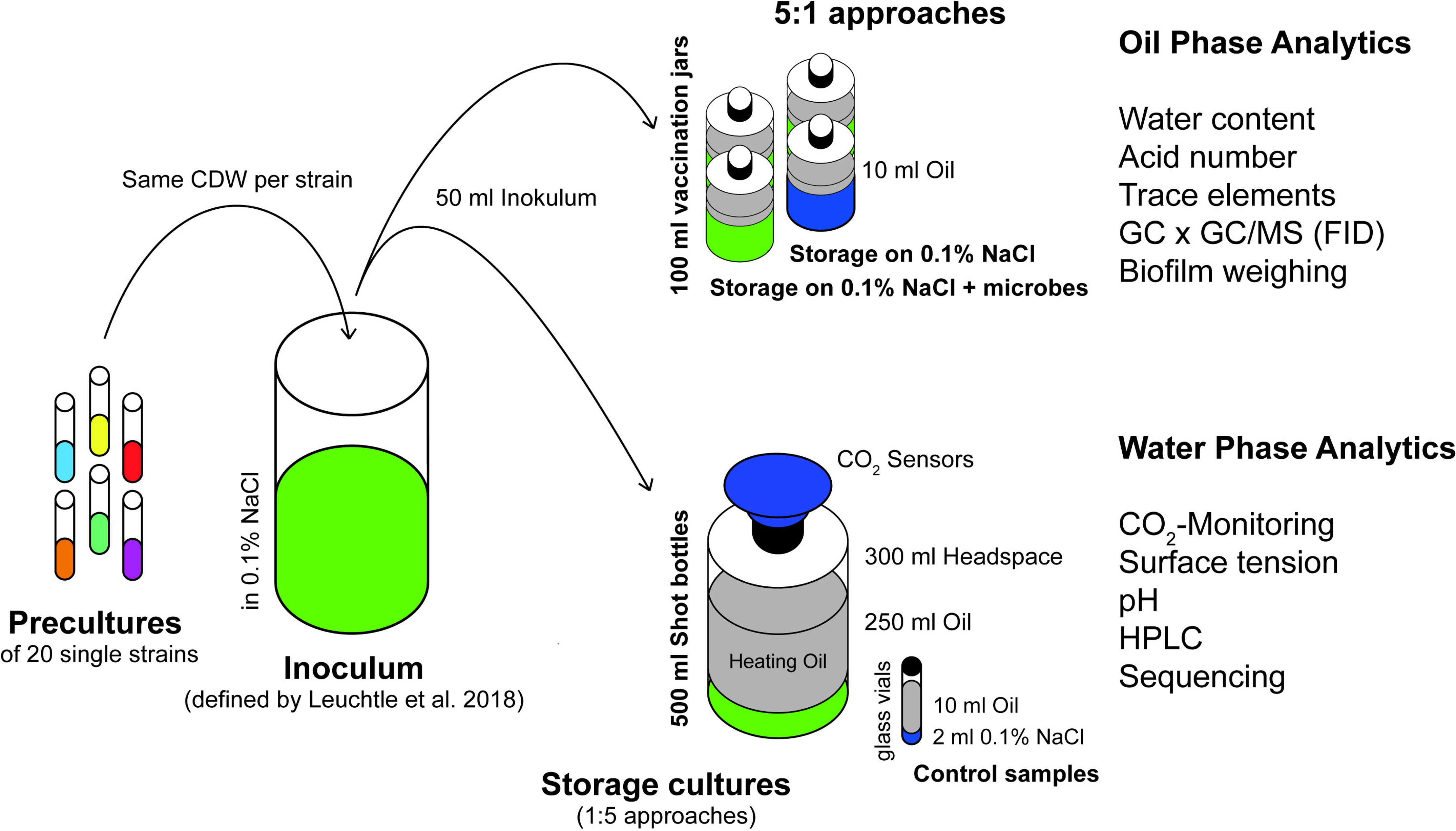
Overview of used storage cultures. Shown are the two different types of storage cultures, used in this study, and the setup of those storage cultures using a defined inoculum of twenty strains (Table 2), representative of heating oil contamination, as defined by Leuchtle et al. (2018) [8]. The classical storage cultures (1:5 approaches) are intended for the CO_2_ measurement and water phase analytics (surface tension, pH, HPLC, sequencing), and the alternative storage cultures (5:1 approaches) are meant for oil phase analytics (water content, acid number, trace elements, GC x GC/MS or GC x GC/FID) and biofilm quantification.

**Table 2.** Strains used in this study and for the defined inoculum of the heating oil tank simulation [8].

| Strain | Database number | Source |
| --- | --- | --- |
| *Acinetobacter beijernickii* | DSM 22901 | DSMZ |
| *Acinetobacter venetianus* | DSM 23050 | DSMZ |
| *Burkholderia cepacia* | DSM 7288 | DSMZ |
| *Burkholderia xenovorans* | - | Leuchtle et al. (2018) |
| *Micrococcus luteus* | - | Leuchtle et al. (2018) |
| *Micrococcus yunnanensis* | DSM 21948 | DSMZ |
| *Pseudomonas fluorescens* | - | Leuchtle et al. (2018) |
| *Pseudomonas poae* | - | Leuchtle et al. (2018) |
| *Candida cylindracea* | DSM 2031 | DSMZ |
| *Debaryomyces hansenii* | DSM 70244 | DSMZ |
| *Debaryomyces polymorphus* | DSM 70816 | DSMZ |
| *Pichia membranifaciens* | DSM 21959 | DSMZ |
| *Raffaelea sp.* | - | Leuchtle et al. (2018) |
| *Rhodotorula mucilaginosa* | DSM 18184 | DSMZ |
| *Ustilago maydis* | - | Leuchtle et al. (2018) |
| *Yarrowia deformans* | CBS 2071 | CBS-KNAW |
| *Yarrowia lipolytica* | - | Leuchtle et al. (2018) |
| *Paecilomyces lilacinus* | DSM 846 | DSMZ |
| *Penicillium chrysogenum* | DSM 21171 | DSMZ |
| *Penicillium citrinum* | - | Leuchtle et al. (2018) |

### 2.2 CO_2_ measurement

For the measurement of CO_2_ development, we used the BCP-CO_2_ system (BlueSens Gas Sensor GmbH). The CO_2_ concentration is monitored by a source of infra-red light, which is weakened by the analyte gas and reflected into the detector unit of the sensor. The sensor was attached airtight to the opening of a culture vessel, which was in this study a 1 L shot bottle. Measurements were performed without air exchange up to two weeks. Oxygen availability of microbial metabolism limits longer runtimes.

### 2.3 Surface tension and pH measurements

The Force Tensiometer K11 (KRÜSS) and a pH meter (Hanna Instruments) were used to measure the surface tension and pH of the separate water phases in the storage cultures. 500 µ l portions of sterile filtered water phase were used.

### 2.4 High-performance liquid chromatography (HPLC) measurements

Samples of the free water phase were harvested after one week and analyzed by HPLC without further treatment. The control samples consisted of 2 ml 0.1 % NaCl without microbes under 10 ml of fuel in 12 ml glass vials. An UltiMate 3000 series instrument from Dionex-Themo Scientific was used for HPLC-RI analysis. A Metab-AAC HPLC column of the BF series from ISERA (300 x 8 mm), intended for the analysis of sugars, organic acids, and alcohols, was used. 5 mM H_2_SO_4_ was applied as running buffer at a flow rate of 0.6 ml/min. The column oven was kept at 75°C. The alcohols were detected via the IR detector unit Refracto Max 521 (running at 35°C with a data collection rate of 10 Hz and an integrator range of 500 µ RIU/V). Quantification was based on peak areas and occurred via an external standard curve of the compound.

### 2.5 Gas-Chromatography (GC-FID) for oxymethylene ether (OME) solutions

A Trace-5 gas chromatograph from Thermo Fisher with a VF-WAXms column (60 m / 0.25 mm / 0.25 µ m) from Agilent was used for GC analysis. Helium was applied as carrier gas at a pressure of 1.3 bar. The injector and FID detector were operated at 250°C. The following temperature program was run in the column oven: 10 min. at 50°C, with 8°C/min. to 250°C, 30 min. at 250°C. The mass recovery was calculated as the difference between the mass of the total sample weighed in and the mass of the OME components found and quantified in the sample by GC-FID. Quantification is performed by peak area comparison with the 1-pentanol standard used and taking into account pre-calculated correction factors for the individual OME components. The GC method does not take into account the OME component OME-6.

### 2.6 Determination of the water content in oil phases

The water content was determined in accordance with DIN EN ISO 12937 via coulometric titration as per Karl Fischer for mineral oil products with boiling points below 390°C [103].

### 2.7 Determination of the acid number in oil phases

The acid number was determined according to EN 14104 “Products of vegetable and animal fats and oils - Fatty acid methyl esters (FAME) - Determination of acid number” [104].

### 2.8 Trace element analysis in oil phases

A screening of trace elements in oil phases was carried out as a contract analysis at ASG Analytik-Service AG using mass spectrometry with inductively coupled plasma (ICP-MS).

### 2.9 Determination of the nitrogen content in oil phases

The determination of the nitrogen content in oil phases was carried out as a contract analysis at ASG Analytik-Service AG using the combustion method with a chemiluminescence detector according to DIN 51444 [105].

### 2.10 Extraction of genomic DNA from storage cultures

For the isolation of genomic DNA from water phases of storage cultures or the inoculum used for storage cultures, a mechanical extraction procedure in combination with chemical purification was chosen to achieve minimal discrimination of individual bacterial, yeast, or mold strains. When extracting from water phases of completed storage cultures, always the complete water phase was collected and extracted as one sample, i.e. including biofilm of the interface and the sediment.

The cell suspension was centrifuged in a 50 ml reaction tube (10 min, 12,000 rpm, 4°C). The supernatant was removed by suction. The remaining supernatant including the cell pellet was frozen overnight at -20°C. The frozen remaining supernatant was removed using a SCANVAC CoolSafe bench top freeze dryer (Labogene) overnight and the cell pellet was completely dried. The cell pellet was resuspended in 1.2 ml of lysis buffer (2% Triton X-100, 1% SDS, 100 mM NaCl, 1 mM EDTA, 10 mM Tris-HCl pH 8.2). The cell suspension was divided into two 2 ml screw cap reaction tubes (600µl each) (MP Biomedicals) filled with 0.7 g ceramic beads (soilGEN, diameter 125 µ m - 250 µ m). For mechanical disruption, the FastPrep 24 5G homogenizer (MP Biomedicals) was used with the following program: 4 m/s, 3 cycles of 30 s each (stored on ice for 30 s in between). The reaction tubes were then stored on ice and centrifuged at 14000 rpm, 4°C until the foam settled completely. The supernatant (approximately 600 µl) was transferred to new 2 ml reaction tubes. 100 µ l NaCl (5 M) and 80 µ l CTAB (10%(w/v) in 0.7 M NaCl) heated to 37°C was added to each tube and mixed thoroughly. The suspension was incubated at 65°C, 500 rpm for 10 min. 600 µ l of phenol:chloroform:isoamyl alcohol (25:24:1) was added. The mixture was shaken for at least 30 s and then centrifuged at 12,000 rpm, 25°C for 5 min. The upper phase was transferred to a new 2 ml reaction tube. 600 µl of chloroform:isoamyl alcohol (49:1) was added and shaken thoroughly. The suspension was centrifuged at 12,000 rpm, 25°C for 5 min and the upper phase was transferred to a new 1.5 ml reaction tube. 500 µ l of isopropanol (ice cold) was added and DNA was precipitated by inverting 30 times. After centrifugation for 10 min at 12,000 rpm, 25°C the supernatant was removed and 700 µl of 70%(v/v) ethanol (cold) was added to the pellet. The mixture was incubated at 25°C for 15 min and then centrifuged at12,000 rpm, 25°C for 5 min. The supernatant was removed as far as possible and the DNA pellet was air dried. The pellet was resuspended in 25 µ l Tris-HCl (10 mM, pH 8.0) overnight at 37°C, 500 rpm. The mixtures of one culture sample, split before cell disruption, were recombined and stored at 4°C.

### 2.11 Sequencing and bioinformatics analysis

Libraries for Illumina sequencing were prepared with the Illumina DNA Prep (M) Tagmentation kit, using 100 ng of DNA as input. Sequencing was performed in a 2 x 151 bp mode on a NextSeq 550 using a 300 cycles reagent with unique dual indexing (IDT for Illumina DNA/RNA UD Indexes, Tagmentation). Library preparation and sequencing were done by the Institute of Medical Microbiology and Hygiene, NCCT Microbiology (Tübingen, DE).

The raw Illumina data conversion and demultiplexing were performed with the <u>nxf-bcl</u> pipeline [106]. The raw reads were first filtered and trimmed using the <u>nxf-fastqc</u> pipeline [107], which uses the <u>fastp</u> program with default filter and trimming parameters. The filtered reads were then mapped with minimap2 [107] against a custom database that contains the genomes from the microbial mix used as inoculum. The alignment PAF files were filtered to retain only full-high-qualityigh quality alignments (> 120 residues matching per read and mapping quality ≥ 50). The alignment files were used to extract summary information about the number of reads mapping to each species. The data processing was done by Dr. rer. nat. Angel Angelov (Freising, DE).

From reads per species per sample the shares of species per sample were calculated. The fold change of the shares was calculated between samples taken at the end of the storage cultures and samples of the inoculum used to start the storage cultures.

### 2.12 Identification/Semi-quantification by GCxGC/MS and quantification by GCxGC/FID of oil phases

Before injection 25-30 mg of the oil phase was dissolved in 20 ml of n-pentane and 10 µ l of cholestane (2 g/L) or octanoic acid ethyl ester (C8:0-Et, 2 g/L) was added as internal standard.

Analysis was done using a GCxGC system (Thermo Scientific, JEOL, and Zoex) of Brechbühler AG (Schlieren, Switzerland) [108]. 1 µ l sample was injected using a PTV on-column injector with an untreated fused silica precolumn (0,5 m x 0,53 mm ID). Helium was applied as carrier gas with 1 ml/min. The compounds were eluted on a polar DB-17 HT column (15 m x 0.25 mm ID, 0.15 µ m film) in the 1^st^ dimension and on a non-polar PS-255 column (3 m x 0.15 mm ID, 0.04 µm film) in the 2^nd^ dimension. Per column, the following program was used: 3 min. at 36 °C, 5 °C/min. to 320 °C. Before the second column, the compounds were trapped in a loop by a cold jet and released after a modulation time of 7.0-7.5hot jet hot-jet. The second column was followed by a time of flight mass spectrometer (ToF-MS) for identification and semi-quantification with a separation range of 40-800 amu and working at 50 Hz. Alternatively, the second column was connected to a flame ionization detector (FID) working at 350 °C with 350 ml/min. air, 35 ml/min. H_2_, and 30 ml/min. N_2_. The measurement was done by Laboratory Lommatzsch & Säger (Cologne, Germany).

### 2.13 Identification/Relative quantification by GC-MS/MS of water accommodated fractions

#### Extraction of water accommodated fractions (WAFs) of fossil heating oils

In a separating funnel, 80 mL of sterilized 0.1 % NaCl solution were carefully overlaid with 0.5 ml of the respective heating oil, to prevent emulsions. After 72 h, 30 mL of WAF were extracted with 3mL of cyclohexane by shaking for 30 min at 100 rpm and 25°C. The WAF extracts were transferred into GC vials and stored at -24 °C until analysis. Solvent controls of cyclohexane were sampled and stored accordingly.

#### GC-MS/MS analysis of WAFs

The cyclohexane extracts of WAFs were analyzed by GC-MS/MS using a 7890A GC coupled with a 5975C triple-quadrupole-MS from Agilent Technologies.the 1 µ L of sample was injected into the GC inlet and heated according to the following oven temperature program: 3 min. at 45 °C, 3 °C/min. to 210 °C, 2 min. atvaporizedThe vaporised compounds were separated in a nonpolar-low polar OPTIMA 5-MS Accent (30 m x 250 µ m x 0.25 µ m) fused silica, capillary column coated with a silarylene stationary phase (MACHEREY-NAGEL GmbH & Co., Germany). Helium was applied as carrier gas with 1.5 ml/min and a pressure of 12.319 psi (0.8494 bar). The sample transfer line into the MS was heated to 220 °C.

The compounds were eluted into the triple-quadrupole-MS, ionized (precursor ions), fragmented by collision with helium gas, and analysed based on the resulting mass spectra of fragmentation ions. Scan mode MS analysis obtained the entire mass-to-ratio (m/z) range for each compound for identification.

#### Identification/Relative quantification of hydrocarbon compounds

Peak detection and integration were done in Agilent G1701EA MSD Productivity ChemStation (Agilent Technologies) using the ChemStation Integrator and the following integration parameters to increase detection sensitivity: initial area reject: 0, initial peak width: 0.046 min., shoulder detection: off, initial threshold 14.0.

For identification, the retrieved mass spectra were matched to compounds registered in the NIST 11 mass spectral database (National Institute of Standards and Technology, USA) using the probability-based matching (PBM) algorithm. The following PBM search parameters were applied as recommended by the manufacturer: significance (uniqueness U and abundance A) U+A = 2, tilting= on, flag threshold= 3, cross-correlation sort= off, minimum estimated purity= 50 %, low report MW=0, high report MW= 9999, maximum hits= 20, remove duplicate CAS numbers= off. The PBM search produced probability-ranked match reliabilities. As stated by the manufacturer, match probabilities of less than 50 % imply substantial differences between the unknown and the reference mass spectra. Therefore, only peaks with match probabilities of 50% or higher were considered as reliably identified, corresponding to 93.81-99.22 % (96.73±1.83 %) of the total peak area detected.

Due to the complexity of the WAF extracts and the large number of identified compounds, absolute quantification using identical analytical standards was not feasible. The analyzed compounds were therefore quantified by the relative percentage of total peak area detected in the sample (peak area [%] = 100*compound peak area/total peak area).

For better comprehensibility, compounds were grouped into classes based on structural similarities, such as aromaticity and degree of alkylation or hydrogenation.

Detectable but small amounts of organosilicon compounds most likely originated from the silicon-based stationary phase of the GC-column due to column bleeding and were therefore excluded from further analyses. To avoid background signals from the solvent in the WAF extracts, peak areas of compounds identified in the solvent control sample were subtracted from the corresponding peak areas found in the WAFs before quantification.

### 2.14 Biofilm weighing

Biofilm weight was calculated by subtracting the weight of empty filters, which were incubated overnight at 100 °C, from the weight of filters loaded with a harvested biofilm that were also incubated overnight at 100 °C. Biofilm was harvested from four biological replicates of 5:1 approaches per oil phase using an inoculation loop. Glass fiber filters with 0.4 µ m diameter were used (GF-5, Macherey Nagel). Weight was measured using a moisture analyzer (MAC 50/1/NH, RADWAG).

### 2.15 Extra light heating oils used in this study

The fossil heating oils (HEL1-6) used in this study represent individual batches from different German refineries, with different crude oil sources and production conditions.

### 2.16 Alternative fuels used in this study

The alternative fuels used in this study are: Rapeseed oil methyl ester, used cooking oil methyl ester, Gas-to-Liquids, hydrogenated vegetable oils, and oxymethylene ethers. They represent individual batches from different German production sites of international companies of the mineral oil or food industry.

## 3. Results

Two culture approaches were used in this work. In the classical storage cultures with 50 ml of water phase and 250 ml of oil phase, therefore also called 5:1 approaches, the effect of the oil phase on the contaminating microbes was investigated. The analyses used were the measurement of microbially produced CO_2_ as a measure of microbial activity (catabolic activity of the biomass present) [37] and water phase analysis. Water phase analysis included surface tension and pH analysis as indicators of the presence of oil phase extracts or microbial metabolites, HPLC measurement to accurately identify and quantify these substances, and sequencing of extracted gDNA to measure microbial diversity. These cultures were performed under bob-permanent aeration and were limited in time to up to three weeks, depending on the available oxygen (Figure 1) [37].

In the alternative storage cultures with 50 ml water phase and 10 ml oil phase, therefore also called 1:5 approaches, the effect of the contaminating microbes on the oil phase was investigated and microbially relevant parameters of the oil phases were determined. These cultures were not subject to oxygen limitation and hence, run longer. Oil phase analysis included water content, acid number, trace elements (plus nitrogen), and GC x GC/MS (FID) (Figure 1).

### 3.1 Fossil heating oils of six different German refineries define the baseline of microbial activity

#### CO_2_-Monitoring (1:5 Ansätze)

To determine differences in microbial susceptibility or possible microbial activity in fossil heating oils, comparative storage cultures of fossil heating oils from six different German refineries (HEL1-6) were performed. For HEL1 and HEL3, microbial activity was found to be 42-126% higher depending on the heating oil used for comparison (Figure 2A).

**Figure 2.**
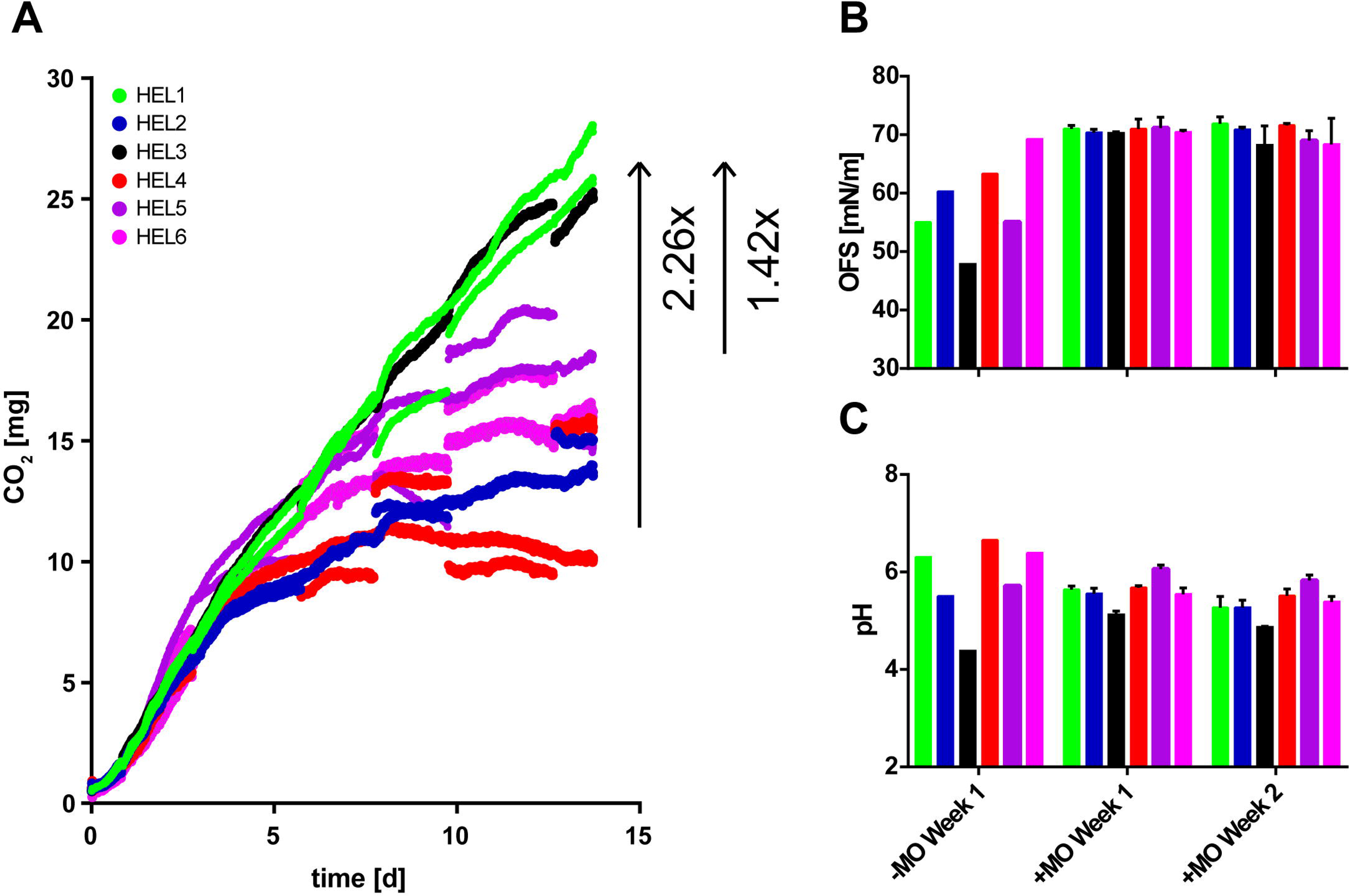
Storage cultures of fossil heating oils. Microbial activity in two-week storage of fossil heating oils from six German refineries measured by the sum of CO_2_ accumulation in oil phase and headspace is shown. Plotted is the discontinuous CO_2_ measurement of three biological replicates (A). The surface tension of free water phases under the tested oil phases after one and two weeks with and without microbes is shown. Plotted are up to three biological replicates (B). The pH of the free water phases after one and two weeks with and without microbes is shown. Plotted are up to three biological replicates (C).

To assess the toxicity or a possible inhibitory effect of the water extract from fossil heating oils on microbes, the water extract of the corresponding heating oils was used as the water phase at the beginning of storage cultures (Figure 3) with heating oils HEL2 and HEL3. The addition of the water extract of HEL2 to a storage culture with HEL2 as an oil phase resulted in a 37% reduction in microbial activity. A comparable inhibition could not be detected for the storage culture with HEL3.

**Figure 3.**
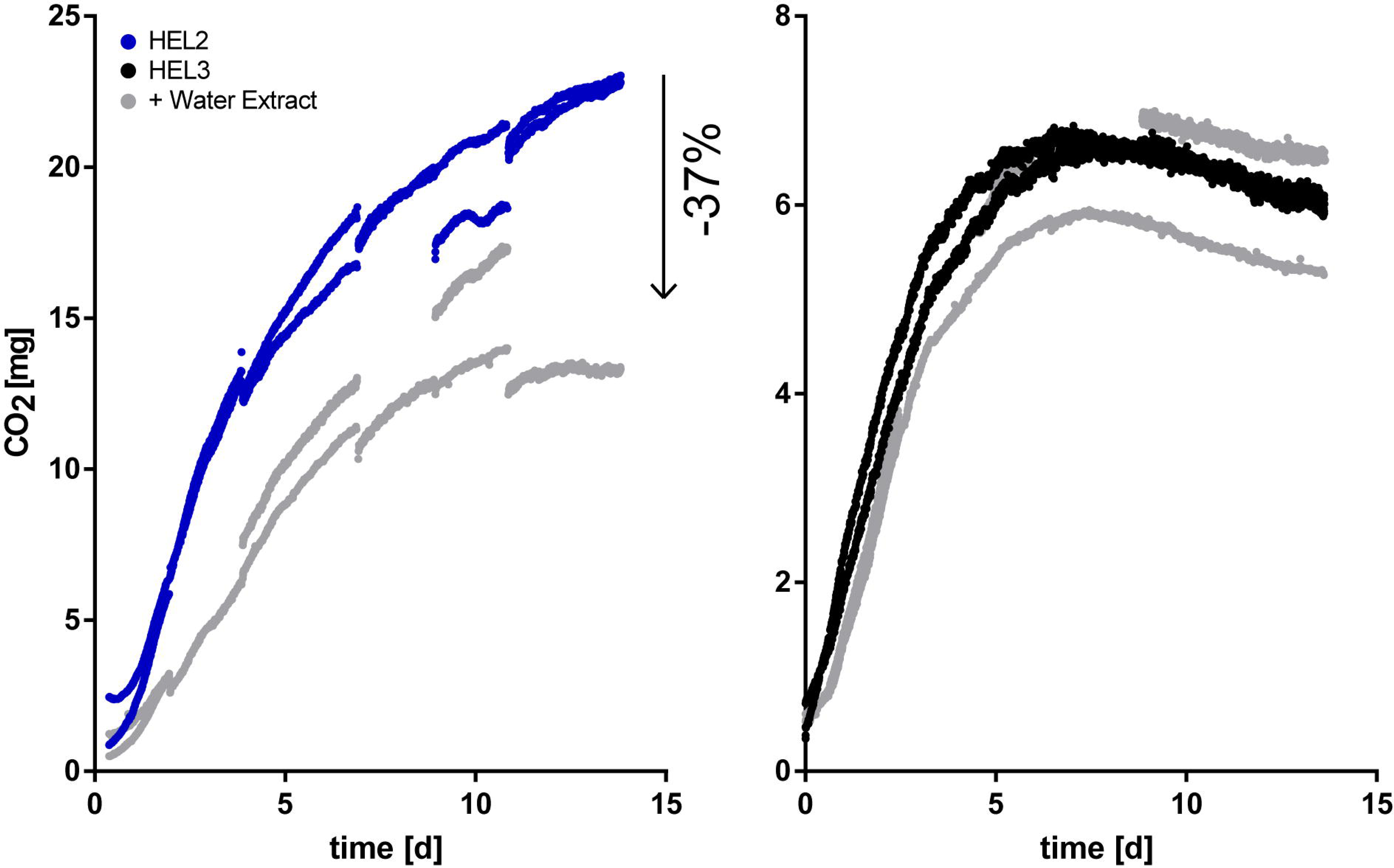
Storage cultures of fossil heating oils HEL2 and HEL3 with and without water extract of the corresponding heating oil as water phase. The microbial activity in a 14-day storage culture with fossil heating oil HEL2 and HEL3 is shown. One time 0.1% NaCl and the other time the water extract of the same heating oil was used as water phase with inoculated microbes. Microbial activity is measured by the sum of CO_2_ accumulation in oil phase and headspace. Plotted is the discontinuous CO_2_ measurement of up to three biological replicates.

Microbial activity is measured here and below as the sum of CO_2_ accumulation in the headspace of the culture bottle and CO_2_ accumulation in the oil phase. Oil phases store a large fraction of the microbially produced CO_2_ and different oil phases store different amounts of the total CO_2_ produced, as shown in previous work [37].

#### Water phase analytics (1:5 Ansätze)

In control samples (water phase without microbes) and the running storage cultures, the pH and surface tension of the water phases were measured after one and two weeks, respectively (Figure 2B and 2C). In the control samples of HEL3, comparatively reduced pH values and reduced surface tensions were observed after one week (-MO Week1), but these were compensated by microbes after one and two weeks (+MO Week 1 and +MO Week 2, respectively).

The GC-MS/MS analysis of the WAF extracts showed that the microbially available fraction of the heating oils is dominated by mono-aromatic compounds (benzenes, xylenes, and cymenes). However, the extracts of the heating oils HEL3, HEL,5 and HEL6 show higher proportions of naphthalenes (di-aromatics) (Figure 4). An assessment of the toxicity of a single fuel or even more a comparison of the toxicity of different fuels requires elaborated microbial toxicity tests, which were not performed as part of the present study. However, a literature search for certain dominant constituents (cymene, xylene, and benzene) shows possible toxic effects on microorganisms or microbial growth-inhibiting effects. Benzene shows growth inhibitory effects on *Pseudomonas aeruginosa* already at 25 µg/L (lowest observed effect concentration; LOEC), whereas *Staphylococcus aureus* shows first effects only at 4 g/L (LOEC) [109]. Cymene on the other hand, appears to be quite non-toxic to microorganisms with a toxicity range of 0.16 - 73 g/L (LOEC; 17 different strains tested) [110–114]. No data have been found for styrene. A comparison to the existing microbial mixture of this study must be taken with caution, however, the literature data show a clear trend regarding a possible toxicity of benzene compared to Cymene. As for the microbial toxicity of di-aromatics compared with mono-aromatics, unfortunately no satisfactory answer can be given here: no data on the microbiological toxicity of relevant di-aromatics (e.g., naphthalenes) were found in the literature search. However, naphthalene is generally regarded the simplest and least toxic of all polycyclic aromatic hydrocarbons (PAHs). Moreover, it has been found, that naphthalene is efficiently degraded at concentrations of 0.5-20 mg/L by *Pseudomonas aeruginosa*, a concentration range where benzene already shows growth inhibitory effects (see above) [115].

**Figure 4.**
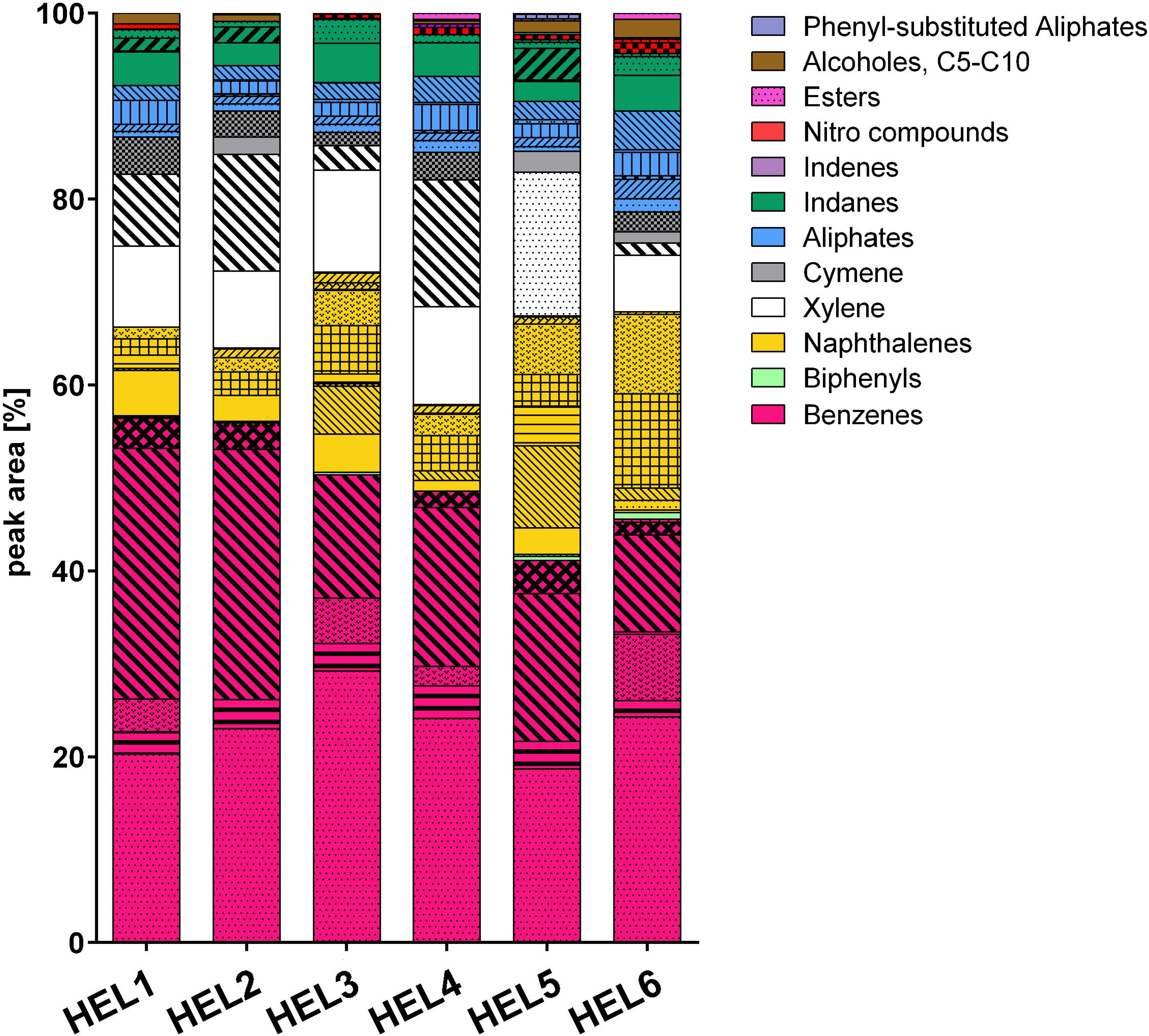
Hydrocarbon composition of water accommodated fractions (WAFs) produced from fossil heating oils (HEL 1-6). WAF extracts in cyclohexane were analyzed by gas chromatography coupled to tandem mass spectrometry (GC-MS/MS). Hydrocarbon compounds in WAF extracts were identified by high matching quality with registered mass spectra (NIST 11 database) and quantified as fraction of the total peak area [%]. Compounds were grouped into classes (colors) and subclasses (patterns) based on structural similarities. The solvent control represents the fraction contributed by the extraction solvent cyclohexane.

The change in reads per strain after completion of storage culture compared to inoculum based on shotgun sequencing showed for all fossil heating oils that bacteria and in particular *Burkholderia* sp. and *Pseudomonas* sp. dominated the culture. However, this bacterial dominance was reduced in the case of the fossil heating oils HEL2, HEL4, HEL5, and HEL6 by more stable proportions of *Y. lipolytica*, *P. penicillatus* (or *P. lilacinus*) and *P. citrinum* (Figure 5 left).

**Figure 5.**
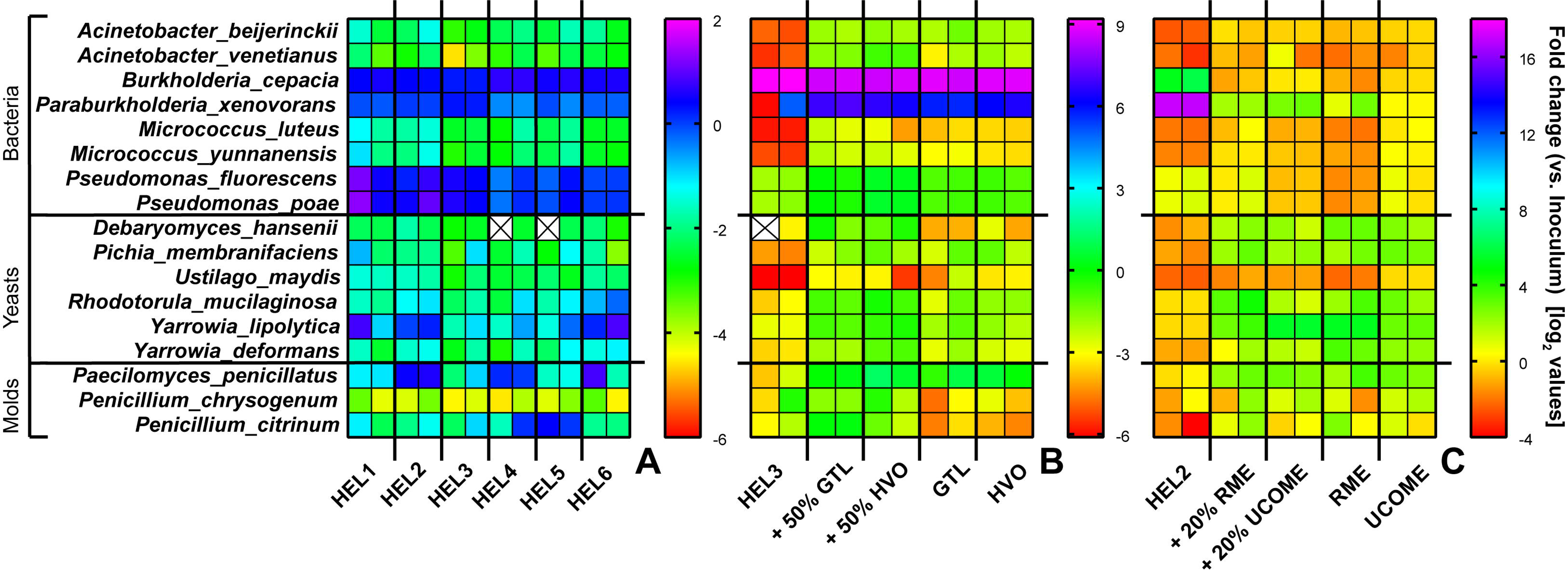
Abundance estimation of microbes by shotgun sequencing following storage cultures. Shown is the result of Illumina sequencing by the Institute of Medical Microbiology and Hygiene, NCCT Microbiology (Tübingen, DE) after cleaning of the Illumina raw data and reads via the “nxf-bcl” and “nxf-fastqc” pipeline and alignment against a user-defined sequence database of the used species by Dr. rer nat. Angel Angelov (Freising, DE). Plotted is the change in reads per species after completion of storage cultures compared to inoculum. A log_2_-scale was applied to the data. Two biological replicates per oil phase are plotted, and the values are mean values of two technical replicates each. The result for the storage culture series of the fossil heating oils is plotted on the left (A), the result for the storage culture series of the GtL/HVO blends is plotted in the middle (B), and the result for the storage culture series of the RME/UCOME blends is plotted on the right (C).

#### Oil phase analytics (5:1 approaches)

Routine analysis attested to higher water contents (30-35 mg/kg versus 20-25 mg/kg) in the untreated heating oils HEL1 and HEL3. Analysis of the water content of the oil phase following two weeks of storage (5 parts water +/- microbes, 1 part oil phase) showed drastic changes. In approaches without microbes, water enrichment of fossil heating oils occurred up to six times. In the presence of microbes, this enrichment is absent (Figure 6 left). To clarify whether the microbes prevent water from entering the oil phase or whether they remove water from the oil phase, an additional experiment was carried out. In this experiment, the storage of fossil heating oil HEL3 on water without microbes for two weeks was followed by two-week storage on water with microbes (Supplementary Figure 1). Here, a water content of approx. 35 mg/kg is measured again after four weeks, as is the case with the untreated heating oil and after only two weeks of storage on water with microbes.

**Figure 6.**
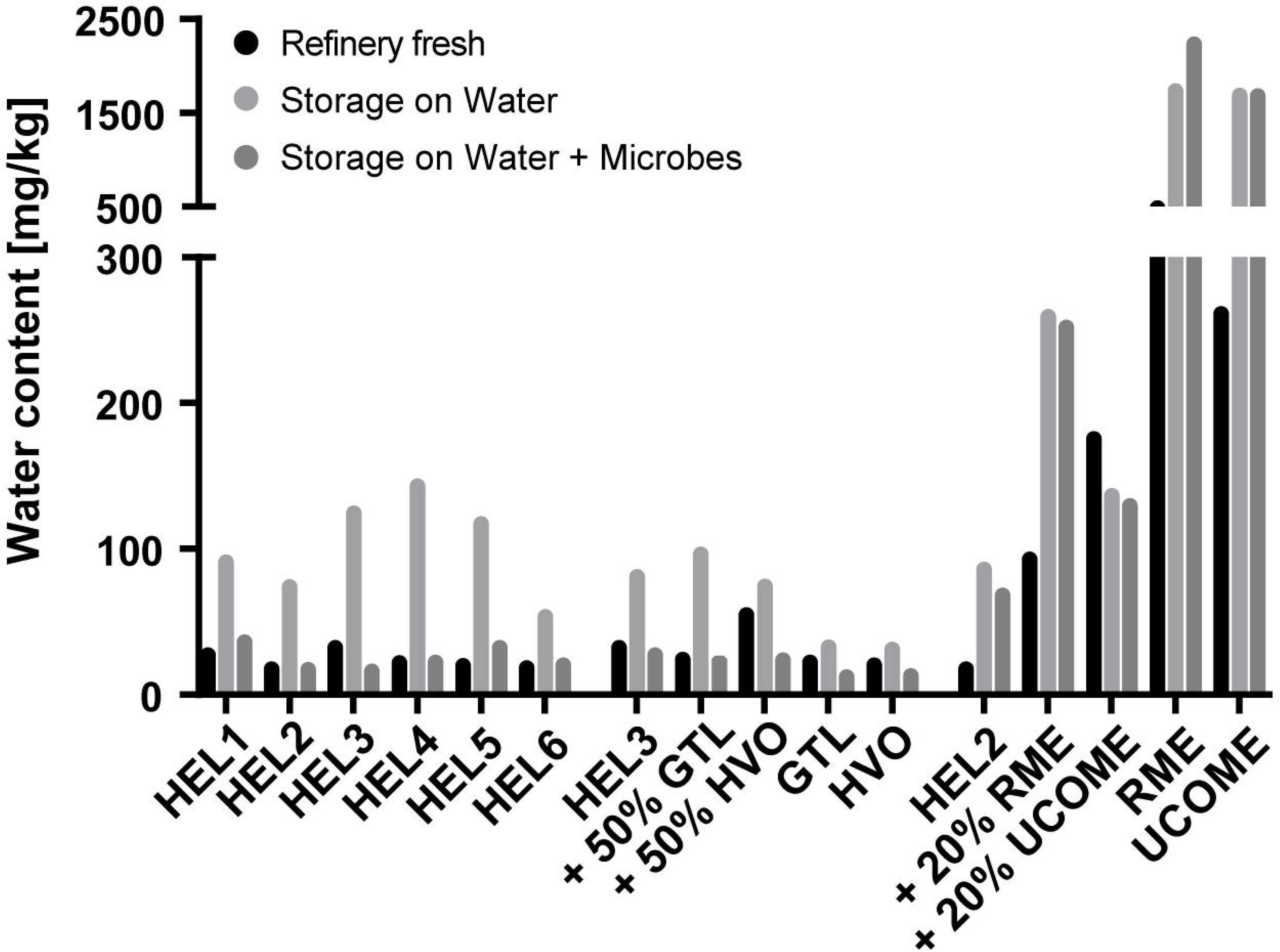
Analysis of the water content in oil phases of 5:1 approaches. The analysis of water content in oil phases of three series of 5:1 approaches is shown. The 5:1 approaches of the fossil heating oils of six German refineries (left), the 5:1 approaches of GTL or HVO blends of the fossil heating oil HEL3 (middle), and the 5:1 approaches of RME or UCOME blends of the fossil heating oil HEL2 (right) are shown. Per oil phase, the water content of the refinery-fresh blend, the water content after two weeks of storage on water without and with microbes are plotted. Shown are the mean values of three technical replicates for refinery- fresh blends or the mean values of technical replicates based on the pooled oil phase of twelve biological replicates when stored on water without and with microbes.

The determination of the trace element content in untreated fossil heating oils was inconspicuous (all measurements below the detection limit of 0.1 mg/kg). However, the fossil heating oils showed different nitrogen contents (Figure 7 top). The heating oils HEL1, HEL3, and HEL5 gave high values of 75.6 mg/kg, 54.1 mg/kg and 141 mg/kg, respectively, while the heating oils HEL2, HEL4 and HEL6 gave comparatively low values of 28.9 mg/kg, 10.6 mg/kg and 15 mg/kg, respectively.

**Figure 7.**
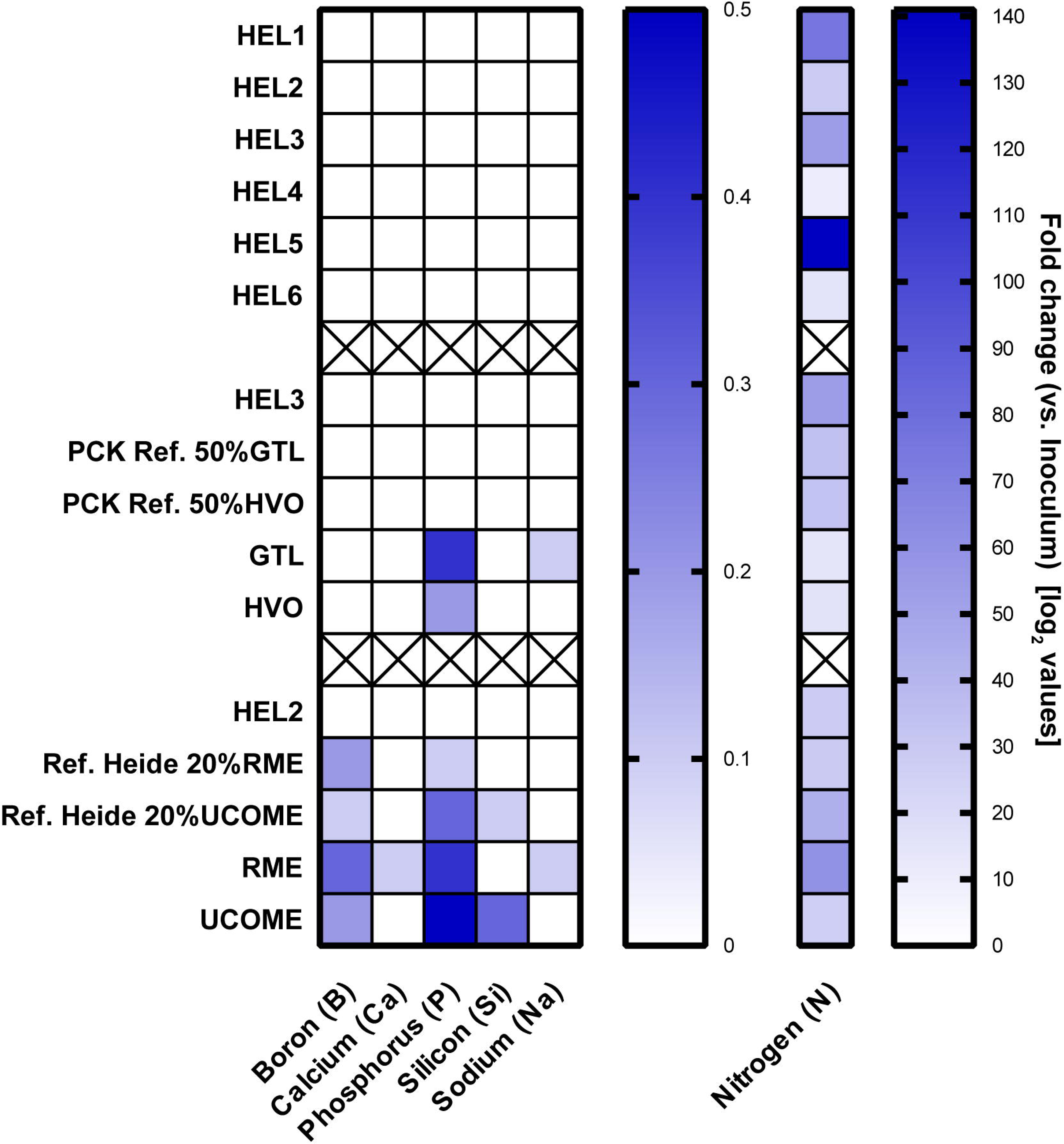
Determination of trace elements and nitrogen contents in refinery-fresh fossil heating oils and alternative blends. The results are ordered according to the storage culture series presented in this paper. Mean values of technical replicates are shown. Trace elements below the detection limit of the test are not shown.

The composition of the heating oils HEL2 and HEL3 was examined in detail by GC x GC/MS (Figure 8). In the case of HEL2, the total ion chromatograms (TICs) show the addition of a paraffin oil consisting of n-/iso- and cyclo-alkanes in the length range C17-C25. In contrast, carbazoles could be detected as possible nitrogen sources in the heating oil HEL3. The exact quantification of the heating oil contents was carried out by GC x GC/FID (Table 10 A and C). With 44% n-/iso alkanes, 34% cyclo alkanes and 22% aromatics, the addition of paraffin oil and the excess of aliphatics were confirmed for the heating oil HEL2. In contrast, the heating oil HEL3 showed an equal distribution of the possible carbon sources with 37% n-/iso alkanes, 30% cyclo alkanes and 33% aromatics. Another difference is the proportion of short-chain aliphatics (C8-C10), which accounts for 9% in HEL2 and only 4% in HEL3. As a result of two weeks of storage, microbial degradation performance was quantified in HEL2 and HEL3 by GC x GC/FID (Table 3 B and D). In the case of HEL2, a total degradation of 26 g/kg was measured, split between 10 g/kg n-/iso alkanes, 6 g/kg cyclo alkanes and 10 g/kg aromatics. For HEL3, a total degradation of 35 g/kg was measured, which is divided into 10 g/kg n-/iso alkanes, 10 g/kg cyclo alkanes and 15 g/kg aromatics.

**Figure 8.**
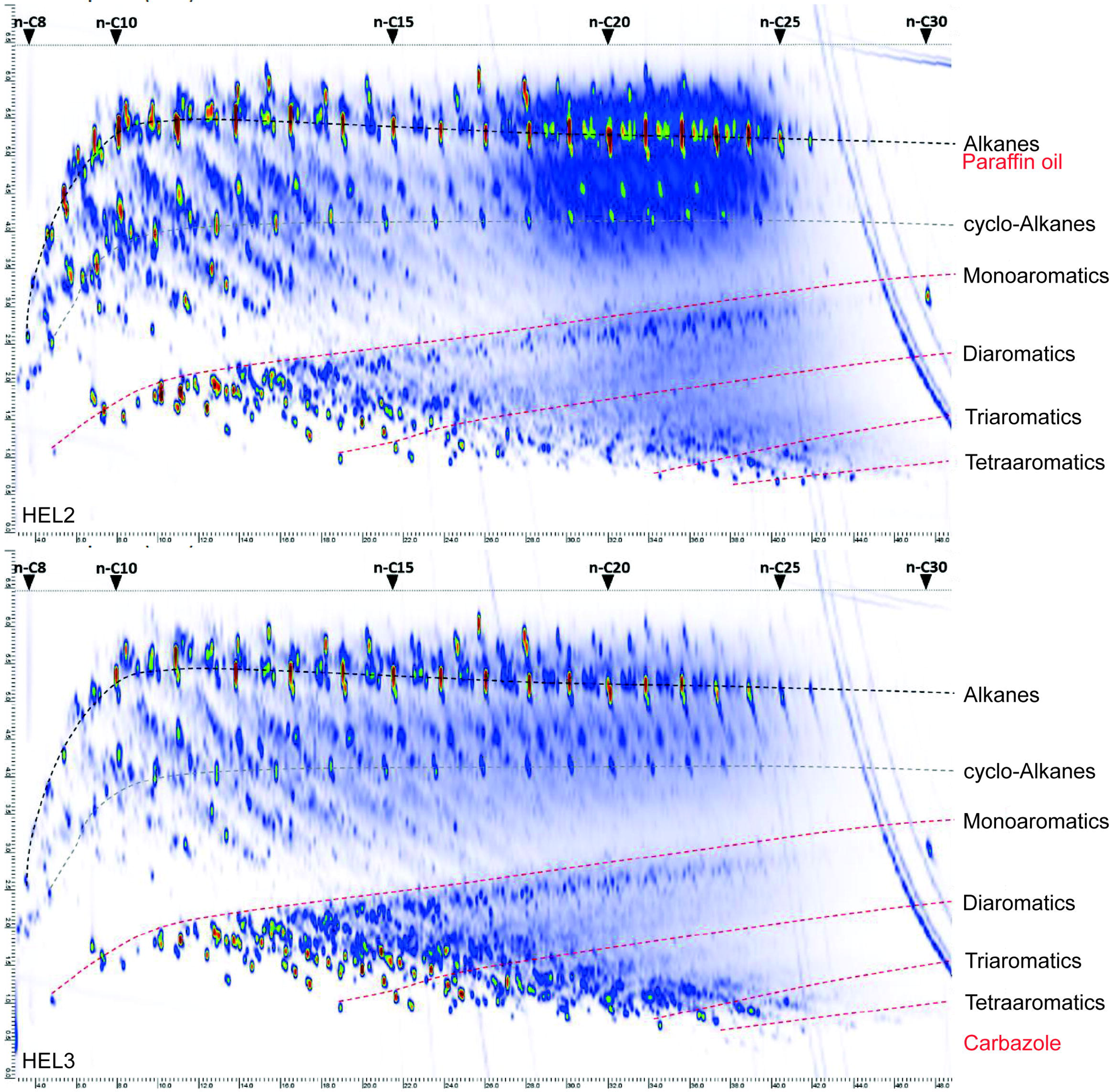
Chemical composition of fossil heating oils HEL2 and HEL3 used in this study. The total ion chromatogram of GC x GC/MS is shown (Laboratory Lommatzsch & Säger, Cologne).

**Table 3.**
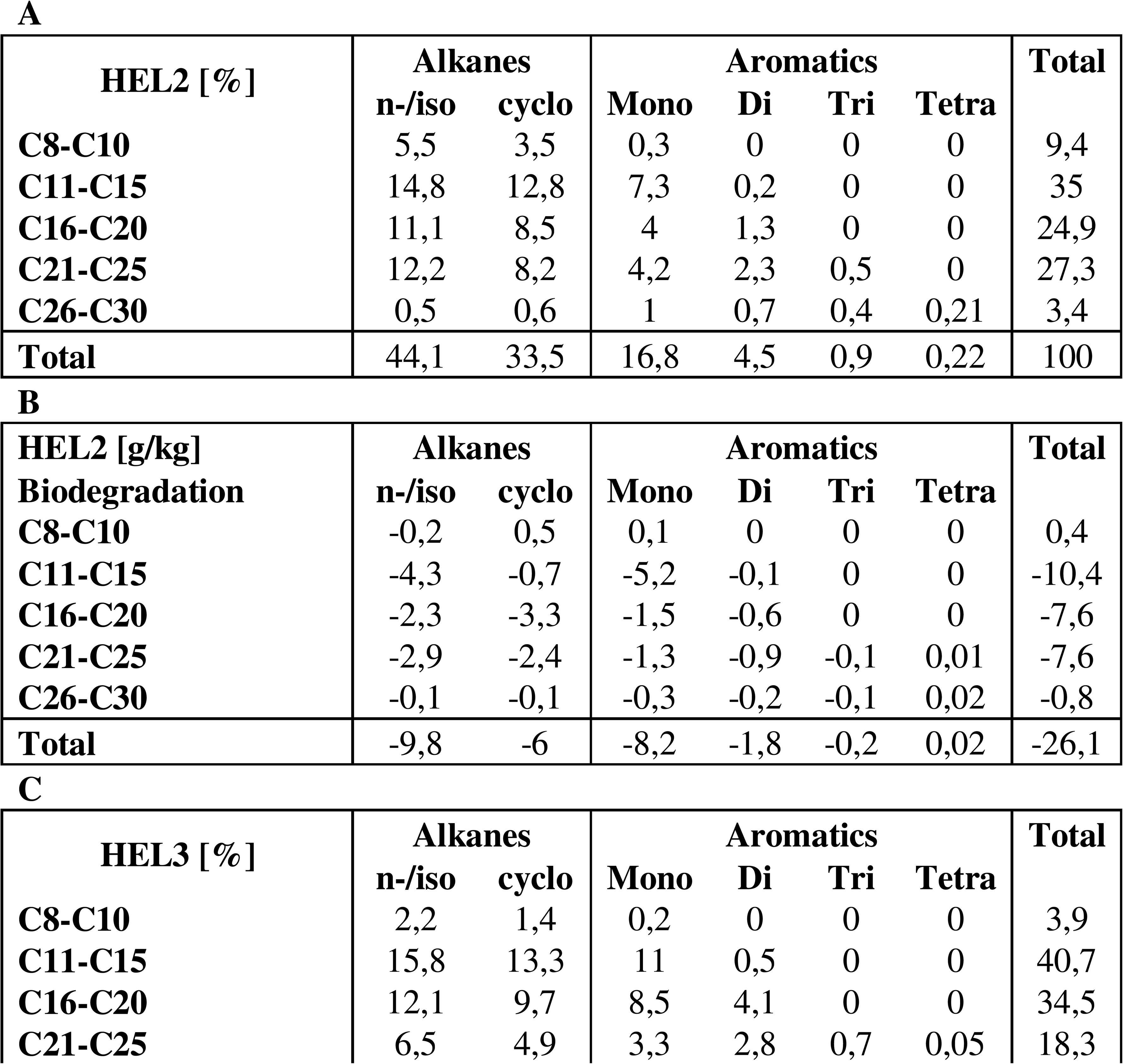
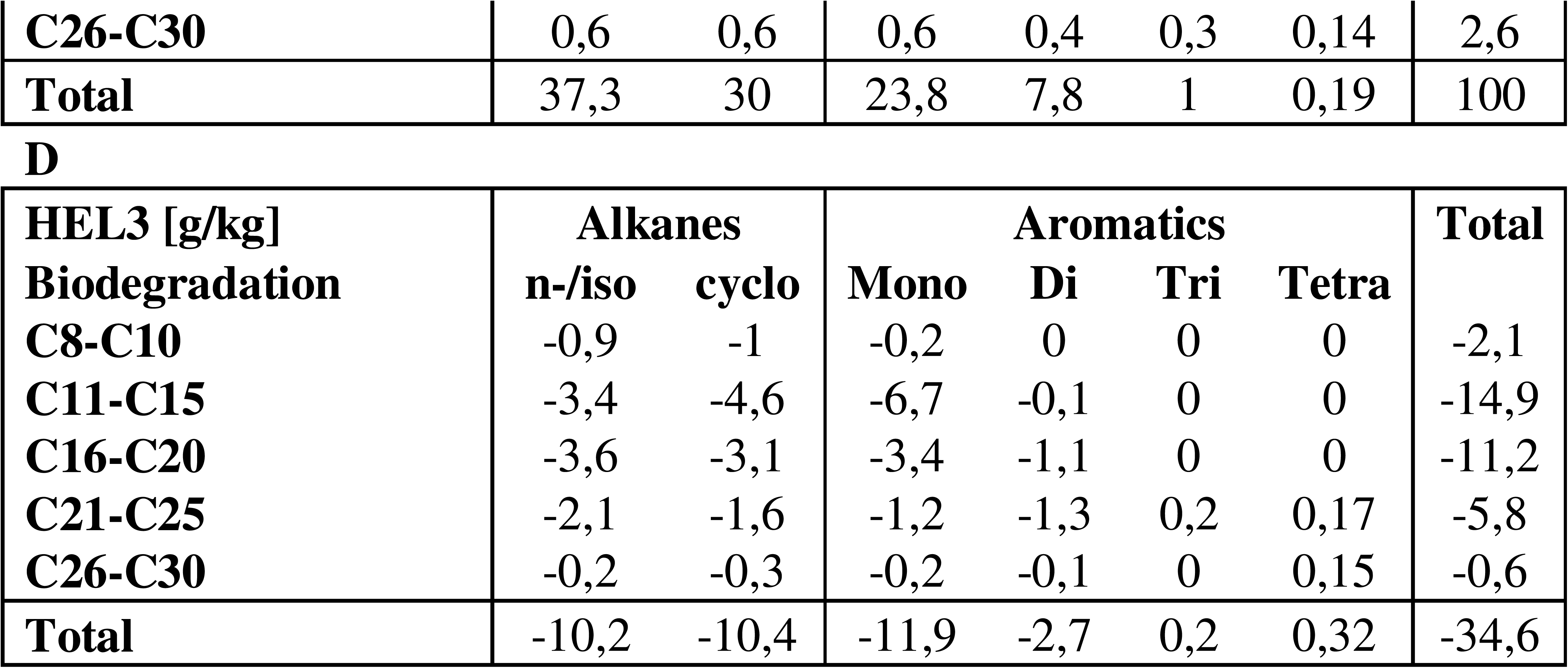
Chemical composition of fossil heating oil HEL2 (A) and HEL3 (C). Degradation of heating oil of HEL2 (B) and HEL3 (D) by microbes during storage cultures in 5:1 approaches. Result of 2D-GC/MS for identification and 2D-GC/FID for absolute quantification (Laboratory Lommatzsch & Säger, Cologne)

| **HEL2 [%]** | **Alkanes** |  | **Aromatics** |  |  |  | **Total** |
| --- | --- | --- | --- | --- | --- | --- | --- |
|  | **n-/iso** | **cyclo** | **Mono** | **Di** | **Tri** | **Tetra** |  |
| **C8-C10** | 5,5 | 3,5 | 0,3 | 0 | 0 | 0 | 9,4 |
| **C11-C15** | 14,8 | 12,8 | 7,3 | 0,2 | 0 | 0 | 35 |
| **C16-C20** | 11,1 | 8,5 | 4 | 1,3 | 0 | 0 | 24,9 |
| **C21-C25** | 12,2 | 8,2 | 4,2 | 2,3 | 0,5 | 0 | 27,3 |
| **C26-C30** | 0,5 | 0,6 | 1 | 0,7 | 0,4 | 0,21 | 3,4 |
| **Total** | 44,1 | 33,5 | 16,8 | 4,5 | 0,9 | 0,22 | 100 |

**B**
| **HEL2 [g/kg]** | **Alkanes** |  | **Aromatics** |  |  |  | **Total** |
| --- | --- | --- | --- | --- | --- | --- | --- |
|  | **n-/iso** | **cyclo** | **Mono** | **Di** | **Tri** | **Tetra** |  |
| **Biodegradation** |  |  |  |  |  |  |  |
| **C8-C10** | -0,2 | 0,5 | 0,1 | 0 | 0 | 0 | 0,4 |
| **C11-C15** | -4,3 | -0,7 | -5,2 | -0,1 | 0 | 0 | -10,4 |
| **C16-C20** | -2,3 | -3,3 | -1,5 | -0,6 | 0 | 0 | -7,6 |
| **C21-C25** | -2,9 | -2,4 | -1,3 | -0,9 | -0,1 | 0,01 | -7,6 |
| **C26-C30** | -0,1 | -0,1 | -0,3 | -0,2 | -0,1 | 0,02 | -0,8 |
| **Total** | -9,8 | -6 | -8,2 | -1,8 | -0,2 | 0,02 | -26,1 |

**C**
| **HEL3 [%]** | **Alkanes** |  | **Aromatics** |  |  |  | **Total** |
| --- | --- | --- | --- | --- | --- | --- | --- |
|  | **n-/iso** | **cyclo** | **Mono** | **Di** | **Tri** | **Tetra** |  |
| **C8-C10** | 2,2 | 1,4 | 0,2 | 0 | 0 | 0 | 3,9 |
| **C11-C15** | 15,8 | 13,3 | 11 | 0,5 | 0 | 0 | 40,7 |
| **C16-C20** | 12,1 | 9,7 | 8,5 | 4,1 | 0 | 0 | 34,5 |
| **C21-C25** | 6,5 | 4,9 | 3,3 | 2,8 | 0,7 | 0,05 | 18,3 |
| **C26-C30** | 0,6 | 0,6 | 0,6 | 0,4 | 0,3 | 0,14 | 2,6 |
| **Total** | 37,3 | 30 | 23,8 | 7,8 | 1 | 0,19 | 100 |

**Table 3. Chemical composition of fossil heating oil HEL2 (A) and HEL3 (C).**
| **HEL3 [g/kg]** | **Alkanes** |  | **Aromatics** |  |  |  | **Total** |
| --- | --- | --- | --- | --- | --- | --- | --- |
| **Biodegradation** | **n-/iso** | **cyclo** | **Mono** | **Di** | **Tri** | **Tetra** |  |
| **C8-C10** | -0,9 | -1 | -0,2 | 0 | 0 | 0 | -2,1 |
| **C11-C15** | -3,4 | -4,6 | -6,7 | -0,1 | 0 | 0 | -14,9 |
| **C16-C20** | -3,6 | -3,1 | -3,4 | -1,1 | 0 | 0 | -11,2 |
| **C21-C25** | -2,1 | -1,6 | -1,2 | -1,3 | 0,2 | 0,17 | -5,8 |
| **C26-C30** | -0,2 | -0,3 | -0,2 | -0,1 | 0 | 0,15 | -0,6 |
| **Total** | -10,2 | -10,4 | -11,9 | -2,7 | 0,2 | 0,32 | -34,6 |

### 3.2 Toxic degradation products of OME blends prevent microbial activity

#### CO_2_-Monitoring (1:5 Ansätze)

During a storage culture of two weeks, the addition of 2% OME to HEL2 led to a reduction of the microbial activity, measured by CO_2_ accumulation. In 550 ml headspace, 34 mg CO_2_ accumulated in the case of pure fossil heating oil, but only 15 mg CO_2_ in the case of the addition of 2% OME, which corresponds to a reduction by 56%. A maximum suppression of microbial activity by 85% was already observed for an admixture of 4% OME (Figure 9A).

**Figure 9.**
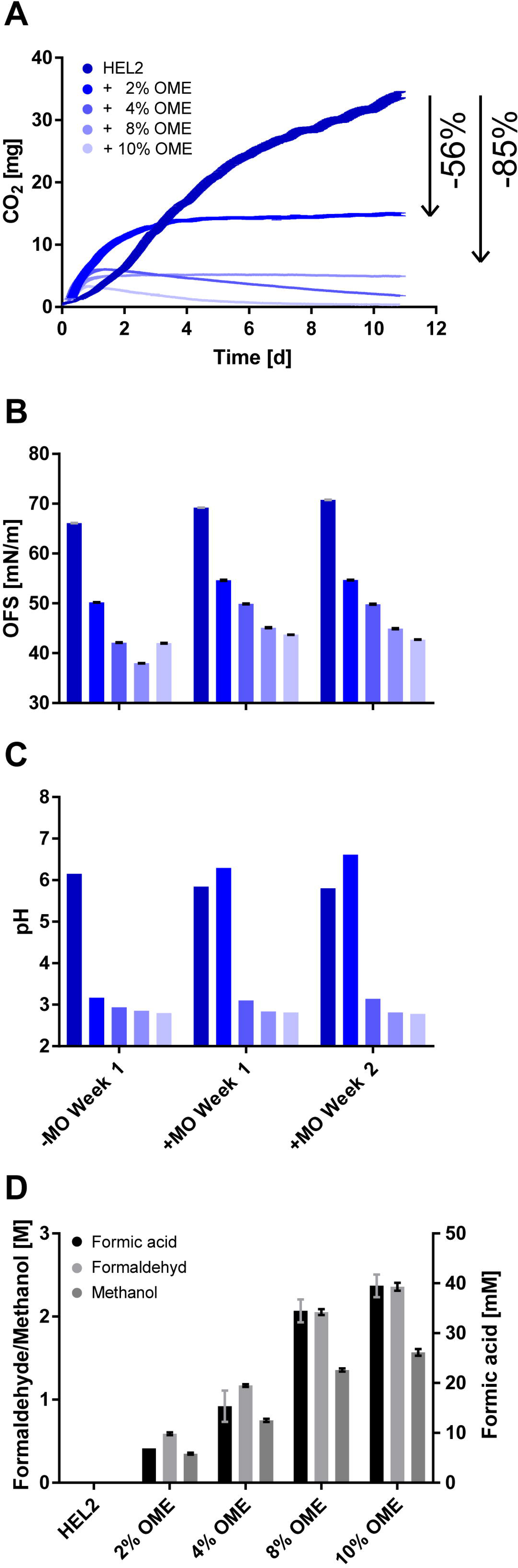
Storage culture of OME admixtures in fossil heating oil. Microbial activity in eleven-day storage of heating oil HEL2 and admixtures of 2-10% OME measured by CO_2_ accumulation in the headspace of the culture bottle is shown. Plotted is the continuous CO_2_ measurement of individual biological replicates (A). The surface tension of free water phases under the tested oil phases after one and two weeks with and without microbes is shown. Plotted are five technical replicates each (B). The pH of the free water phases after one and two weeks with and without microbes is shown. Plotted are individual biological replicates (C). Methanol, formaldehyde, and formic acid concentrations as a result of HPLC analysis of the free water phases after one week without microbes are shown. Two technical replicates each are plotted (D).

#### Water phase analytics (1:5 Ansätze)

The measurement of pH and surface tension of the free water phase, which is carried out in parallel to the CO_2_ monitoring, shows a decreasing surface tension and pH value of the non-buffered water phase with increasing OME content of the oil phase in the presence and absence of microbes. In the presence of microbes, the pH remains neutral at an admixture of 2% OME (Figure 9B and C). HPLC analysis was performed to further identify and quantify the surface-active and acidic substances associated with OME admixture in the free water phase faced by the microbes. In control samples without microbes, formaldehyde was detected at concentrations up to 2.4 M, methanol at concentrations up to 1.6 M, and formic acid at concentrations up to 40 mM after one week, depending on OME admixture (Figure 9D).

#### Oil phase analytics (5:1 Ansätze)

The integrity of the oxymethylene ether solution used for blending (0.2% OME-2; 46.97% OME-3; 29.40% OME-4; 16.66% OME-5; 5.57% OME-6) was confirmed by GC analysis (Supplementary Figure 2).

#### Biomass development in long-term storage (1:5 Ansätze)

To test the sustainability of the antimicrobial effect of the OME admixture, long-term storage of the blend of HEL2 and 10% OME in closed shot bottles was carried out for one year. After one, two and a half, five and twelve months, the developing biomass was measured via quantification of genomic DNA from intact cells. Based on the extractable DNA concentration, a reduction in intact biomass to below 1% occurred within the first month and no re-increase was observed within twelve months (Figure 10).

**Figure 10.**
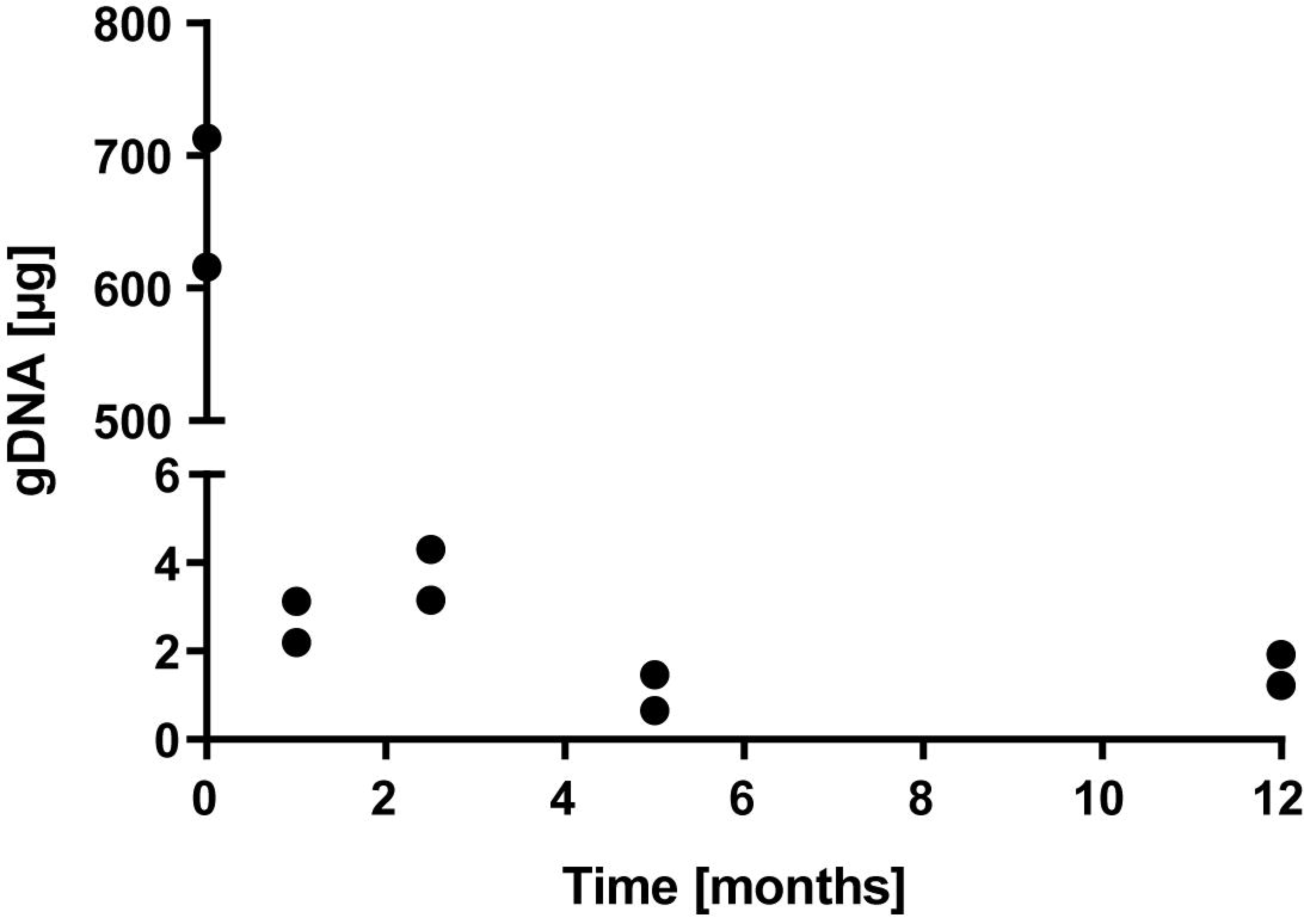
Biomass development in long-term storage of 10% OME blends in fossil heating oil HEL2. As an indicator of biomass, the amount of genomic DNA extractable from whole cells is plotted from each of two biological replicates.

### 3.3 Intensification of the microbial activity below rapeseed oil methyl ester and usedmethyl esterl methylester blends

#### CO_2_-Monitoring (1:5 approaches)

Fossil heating oil HEL2 was chosen as the basis for analyzing the effect of rapeseed oil methyl esters (RME) and used cooking oil methyl esters (UCOME) on possible microbial activity in heating oil storage tanks. The fossil heating oil was compared in an 18-day storage culture with a commercial admixture of 20% RME or 20% UCOME, as well as with oil phases of pure RME and pure UCOME. Under the fossil heating oil, microbial activity, measured by the sum of CO_2_ accumulation in 300 ml headspace and 250 ml oil phase, of about 22 mg CO_2_ was possible. The admixture of 20% RME or 20% UCOME led to an increase in microbial activity of 80-120% to 40-49 mg CO_2_, pure RME or UCOME even allowed an increase of 130-160% to 50-57 mg CO_2_ throughout two and a half weeks (Figure 11A).

**Figure 11.**
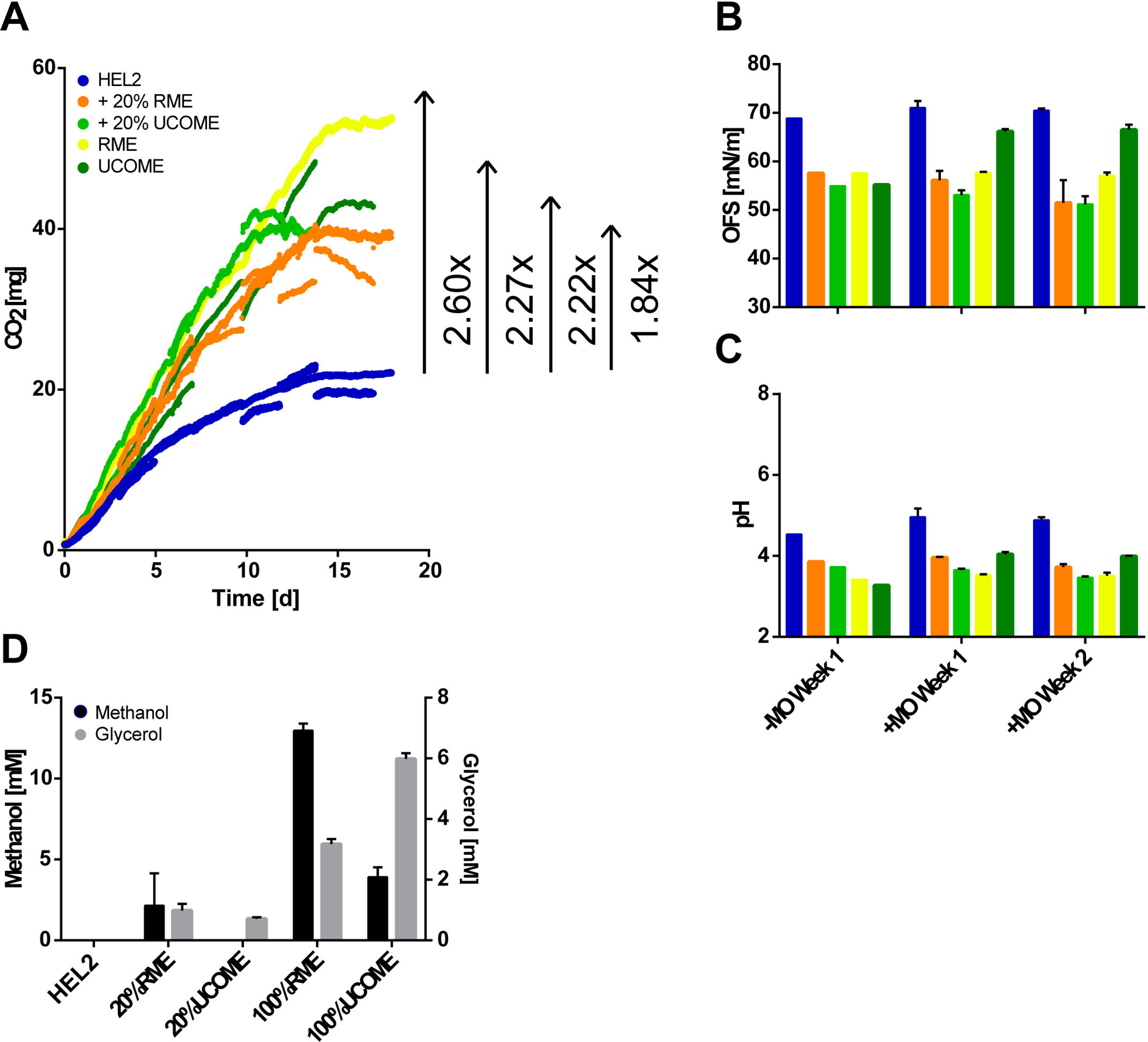
Storage cultures of rapeseed methyl ester (RME) and used cooking oil methyl ester (UCOME) blends. Microbial activity in an 18-day storage culture with fossil heating oil HEL2, fossil heating oil with 20% admixture of RME or UCOME, and pure RME or UCOME is shown. Microbial activity is measured by the sum of CO_2_ accumulation in oil phase and headspace. Plotted is the discontinuous CO_2_ measurement of up to three biological replicates (A). The surface tension of free water phases under the tested oil phases after one and two weeks with and without microbes is shown. Plotted are up to three biological replicates (B). The pH of the free water phases after one and two weeks with and without microbes is shown. Plotted are up to three biological replicates (C). The amount of methanol and glycerol detected by HPLC in the free water phases after one week without microbes is shown. Two technical replicates are plotted (D).

#### Water phase Analytics (1:5 approaches)

Analysis of the water phases showed a reduction in surface tension in the control samples (water phases without microbes) under 20% and 100% RME as well as 20% and 100% UCOME after one week (Figure 11B, -MO Week1), which could only be compensated by microbes in the case of 100% UCOME after one and two weeks (Figure 11B, +MO Week1 and 2). Subsequent HPLC analysis detected glycerol and methanol in the water phases of the control samples (water phases without microbes). Increased methanol (up to 13 mM) was measured in the water phases under RME and increased glycerol (up to 6 mM) under UCOME (Figure 11C). Furthermore, in the water phases of the control samples (water phases without microbes) under 20% and 100% RME, as well as 20% and 100% UCOME, a decrease in pH, was detected after one week (Figure 11D, -MO Week1), which was in no case compensated by microbes after one and two weeks (Figure 11D, +MO Week1 and 2).

The change in reads per strain after completion of the two and a half week storage culture compared to the inoculum based on shotgun sequencing showed a strong increase in the proportions of all yeasts and molds with single exceptions (*Ustilago maydis*) for 20% and 100% RME as well as 20% and 100% UCOME to the same extent. The bacteria show stable proportions or losses except *Burkholderia xenovorans* (Figure 5 right).

#### Oil phase analytics (5:1 approaches)

The oil phase analysis already certified an increase in the water content of the untreated blends 20% RME, 20% UCOME, 100% RME, and 100% UCOME to 96 mg/kg, to 178.45 mg/kg, to 534.15 mg/kg, and to 264.40 mg/kg, respectively, compared to the fossil heating oil HEL2 (20.75 mg/kg). After two weeks of storage of the blends on water, there was a further increase in water content in the case of 20% RME, 100% RME, and 100% UCOME to 262.40 mg/kg, 1781.10 mg/kg, and 1735.55 mg/kg, respectively, compared to HEL2 (88.95 mg/kg). The suppression of water enrichment by storage on water with microbes does not occur (Figure 6 right). The untreated blends of 20% UCOME, 100% RME, and 100% UCOME showed strongly increased acid numbers of 0.786, 0.386, and 0.404 mg KOH per g sample, respectively, compared to HEL2 (0.033 mg KOH per g sample). In the case of 20% UCOME, storage on water with microbes for two weeks resulted in a reduction of the acid number to 0.047 mg KOH per g sample, while in the case of 100% RME and 100% UCOME a further increase to 9.581 and 9.441 mg KOH per g sample, respectively, was recorded (Figure 12 right).

**Figure 12.**
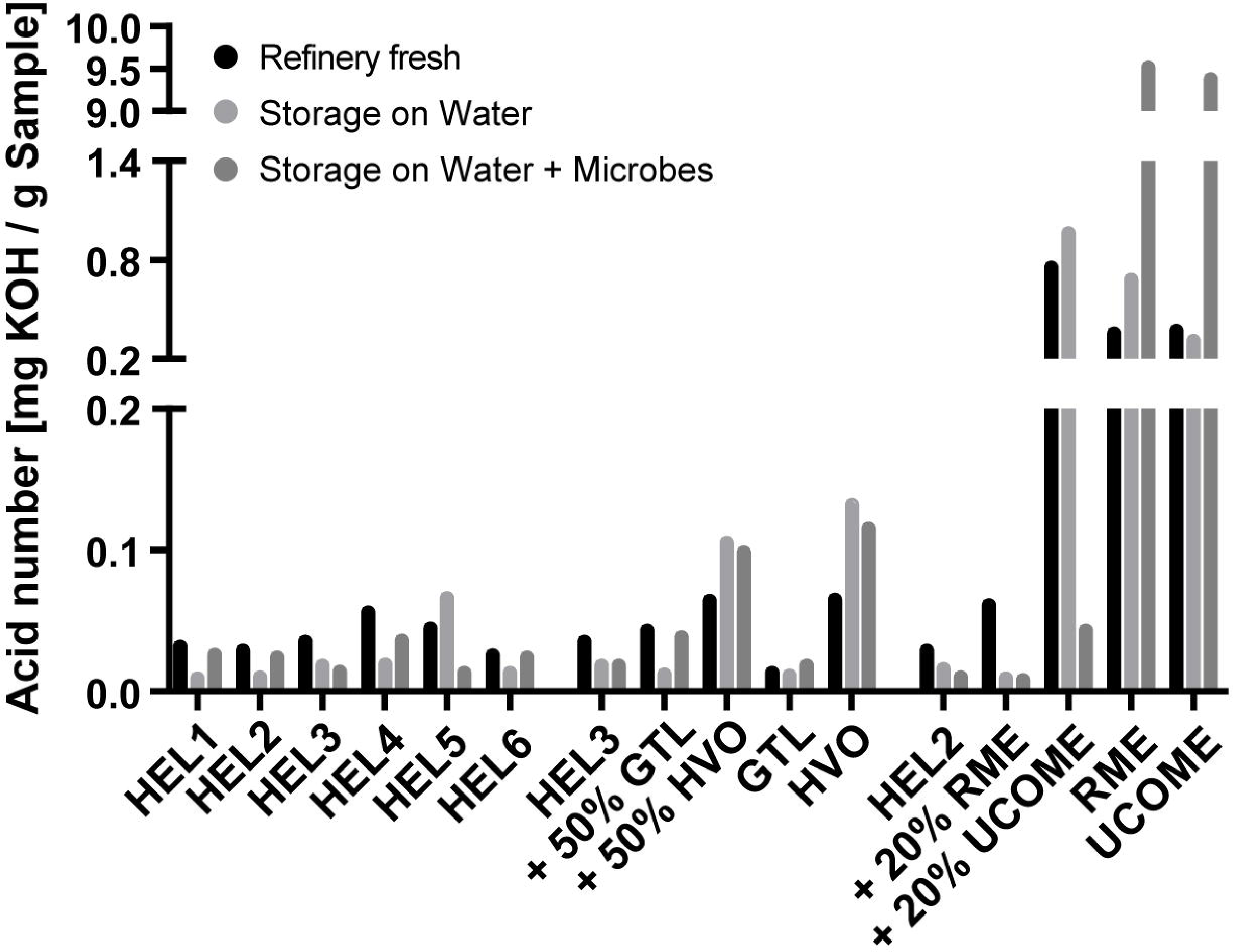
Analysis of the acid number in oil phases of 5:1 approaches. The analysis of acid number in oil phases of three series of 5:1 approaches is shown. The 5:1 approaches of the fossil heating oils of six German refineries (left), the 5:1 approaches of GTL or HVO blends of the fossil heating oil HEL3 (middle), and the 5:1 approaches of RME or UCOME blends of the fossil heating oil HEL2 (right) are shown. Per oil phase, the acid number of the refinery-fresh blend, the acid number after two weeks of storage on water without and with microbes are plotted. Shown are the mean values of three technical replicates for refinery- fresh blends or the mean values of technical replicates based on the pooled oil phase of twelve biological replicates when stored on water without and with microbes.

Trace element analysis of the untreated oil phases attested detectable amounts of boron (0.2-0.3 mg/kg) and phosphorus (0.4-0.5 mg/kg) in the RME and UCOME feedstock. Silicon (0.3 mg/kg) was detectable in UCOME. In addition, the pure RME and UCOME showed significantly elevated nitrogen levels at 60 mg/kg and 26.6 mg/kg, respectively, compared to the fossil heating oil HEL2 at 28.9 mg/kg (Figure 7 bottom).

The composition of RME and UCOME was identified by GC x GC/MS and accurately quantified by GC x GC/FID (Table 4 and Figure 13 bottom). Both RME and UCOME were mainly composed of C18:1 (65.3% and 52.1%, respectively), C18:2 (19.2% and 22.3%, respectively), and C18:3 methyl esters (7.2% and 5.1%, respectively). However, higher proportions of C16:0 (4.3% vs. 11.1%), the additional occurrence of corresponding ethyl esters, and an overall broader chain length spectrum were observed for UCOME (Table 4 bottom).

**Figure 13.**
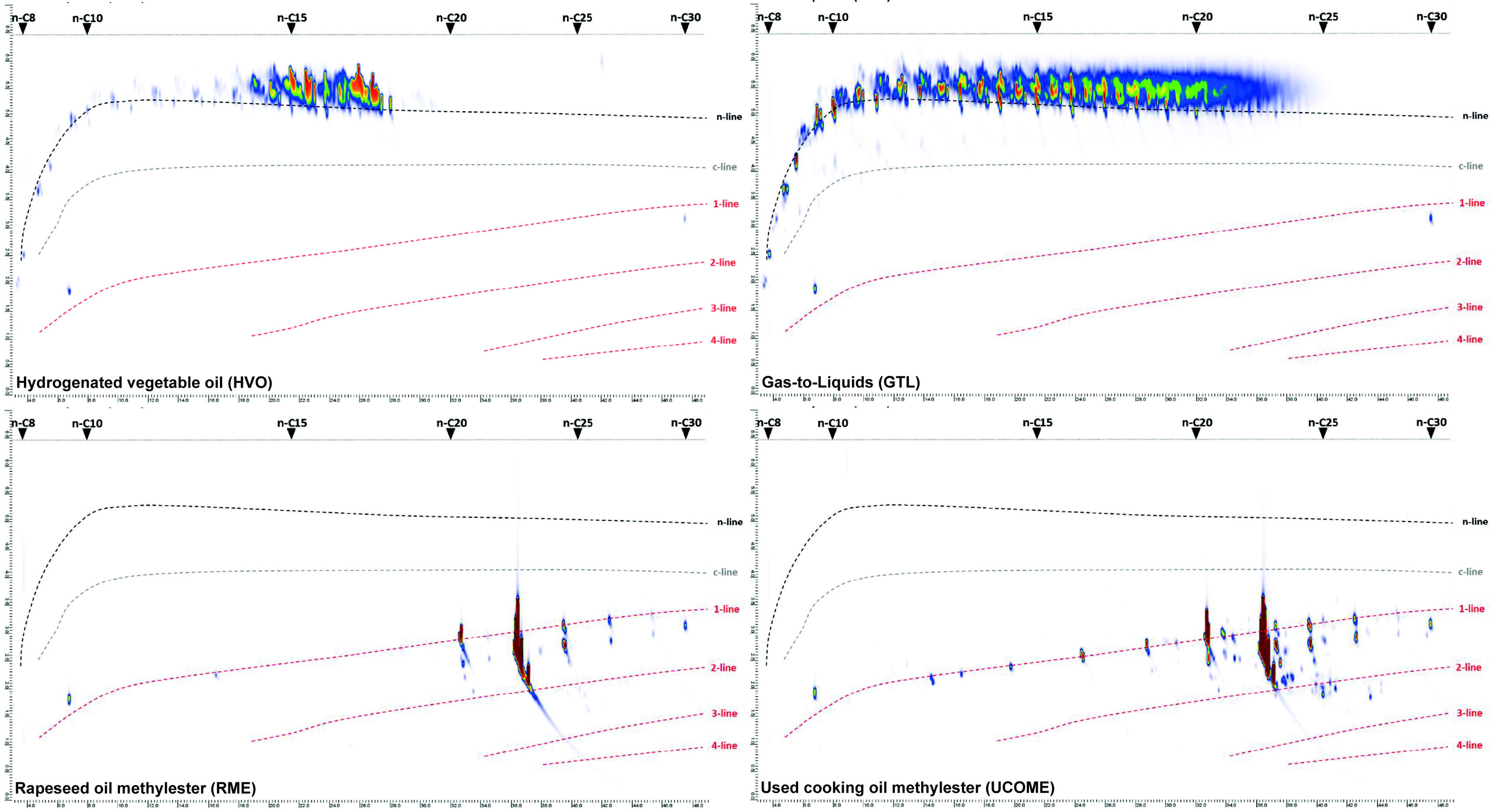
Chemical composition of alternative fuels used in this study. A total ion chromatogram of 2D-GC/MS is shown (Laboratory Lommatzsch & Säger, Cologne).

**Table 4.** Chemical composition of alternative fuels used in this study, results of 2D- GC/MS (Laboratory Lommatzsch & Säger, Cologne)

| **HVO** |  |  | **GTL** |  |  |
| --- | --- | --- | --- | --- | --- |
| **Alkanes** |  |  | **Alkanes** |  |  |
| **[%]** | **n-/iso** | **cyclo** | **[%]** | **n-/iso** | **cyclo** |
| **C8-C10** | 1,2 | 0,1 | **C8-C10** | 7,6 | 0,3 |
| **C11-C15** | 24,2 | 0,2 | **C11-C15** | 38,3 | 1,0 |
| **C16-C20** | 73,1 | 0,6 | **C16-C20** | 38,7 | 0,6 |
| **C21-C25** | 0,2 | 0,0 | **C21-C25** | 13,3 | 0,2 |
| **C26-C30** | 0,3 | 0,0 | **C26-C30** | 0,1 | 0,0 |
| **RME** |  |  | **UCOME** |  |  |
| **Fatty acid** |  |  | **Fatty acid** |  |  |
| **[%]** | **methyl esters** | **ethyl esters** | **[%]** | **methyl esters** | **ethyl esters** |
| **Chain lengths** |  |  | **Chain lengths** |  |  |
| **C8** | 0,0 | 0,0 | **C8** | 0,1 | 0,0 |
| **C10** | 0,0 | 0,0 | **C10** | 0,1 | 0,0 |
| **C12** | 0,0 | 0,0 | **C12** | 0,6 | 0,0 |
| **C14** | 0,0 | 0,0 | **C14** | 0,6 | 0,0 |
| **C16** | 4,8 | 0,0 | **C16** | 11,7 | 0,4 |
| **C17** | 0,1 | 0,0 | **C17** | 0,2 | 0,0 |
| **C18** | 91,9 | 0,0 | **C18** | 80,9 | 1,9 |
| **C19** | 0,1 | 0,0 | **C19** | 0,1 | 0,0 |
| **C20** | 2,3 | 0,0 | **C20** | 2,4 | 0,1 |
| **C22** | 0,6 | 0,0 | **C22** | 0,9 | 0,0 |
| **C24** | 0,2 | 0,0 | **C24** | 0,2 | 0,0 |
| **Double Bonds** |  |  | **Double Bonds** |  |  |
| **0** | 5,4 | 0,0 | **0** | 14,7 | 0,5 |
| **1** | 67,3 | 0,0 | **1** | 54,1 | 1,1 |
| **2** | 19,4 | 0,0 | **2** | 22,5 | 0,3 |
| **3** | 7,3 | 0,0 | **3** | 5,2 | 0,5 |
| **>3** | 0,6 | 0,0 | **>3** | 1,3 | 0,0 |

### 3.4 Limited microbial activity below Gas-to-Liquids and hydrogenated vegetable oils

#### CO_2_-Monitoring (1:5 approaches)

To analyze the effect of gas-to-liquid fuels (GtL) and hydrogenated vegetable oils (HVO) on the possible microbial activity in heating oil storage facilities, the fossil heating oil HEL3 was chosen as a basis. The fossil heating oil was compared in a 17-day storage culture with a commercial admixture of 50% GtL or 50% HVO, as well as with oil phases of pure GtL and pure HVO. Under the fossil heating oil, microbial activity, as measured by the sum of CO_2_ accumulation in 300 ml headspace and 250 ml oil phase, of about 24 mg CO2 was possible.

The admixture of 50% HVO resulted in a 28% reduction of the possible activity to 17 mg CO_2_. The admixture of 50% GtL, pure HVO, and pure GtL allowed even only a microbial activity of 13.3-14.5 mg CO_2_, which corresponds to a reduction of 45% (Figure 14A).

**Figure 14.**
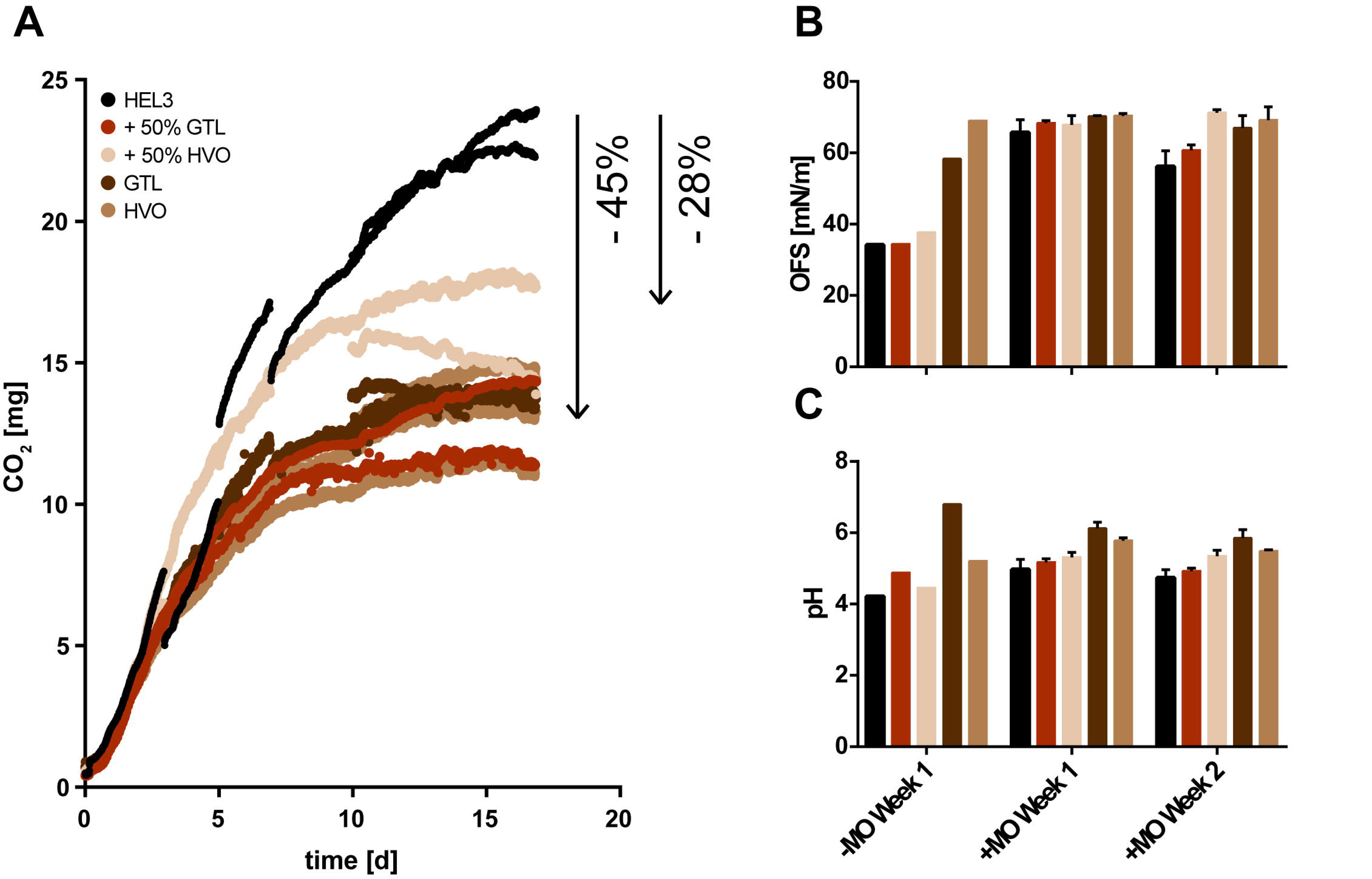
Storage cultures of gas-to-liquids (GtL) and hydrogenated vegetable oil (HVO) blends. Microbial activity in a 17-day storage culture with heating oil HEL3, fossil heating oil with 50% admixture of GtL or HVO, and pure GtL or HVO is shown. Microbial activity is measured by the sum of CO_2_ accumulation in oil phase and headspace. Plotted is the discontinuous CO_2_ measurement of three biological replicates (A). The surface tension of free water phases under the tested oil phases after one and two weeks with and without microbes is shown. Plotted are up to three biological replicates (B). The pH of the free water phases after one and two weeks with and without microbes is shown. Plotted are up to three biological replicates (C).

#### Water phase analytics (1:5 approaches)

In the control samples (water phase without microbes), overlaying with fossil heating oil, with 50% HVO and 50% GtL led to a reduction in surface tension after one week (-MO Week 1), which is absent with pure HVO or pure GtL. This reduction in surface tension is compensated by the presence of microbes after one and two weeks (+MO Week 1 and 2) (Figure 14B). In the water phases of the control samples (water phase without microbes) below pure GtL alone, there was no reduction in pH after one week (-MO Week 1). Reductions in pH in the remaining samples were compensated by the presence of microbes after one and two weeks (+MO Week 1 and 2) (Figure 14C).

The change in reads per strain after completion of the 17-day storage culture compared to the inoculum based on shotgun sequencing showed for 50% GtL and 50% HVO compared to fossil heating oil that bacteria and specifically *Burkholderia sp.* continue to dominate the culture, but also that the majority of yeasts and molds can maintain their shares in the culture (exception *Ustilago maydis*). In contrast, sequencing of samples from storage cultures with pure GtL and pure HVO showed a renewed reduction in the proportions of individual yeasts and molds (*Debaryomyces hansenii*, *Yarrowia deformans*, *Penicillium chrysogenum*, *Penicillium citrinum*) (Figure 5 middle).

#### Oil phase analytics (5:1 approaches)

The oil phase analysis confirmed that the untreated blends of 50% GtL or 50% HVO with the fossil heating oil and the pure GtL or HVO oil phases reduced the increased water content in the fossil heating oil HEL3 from 35 mg/kg to 23-27 mg/kg. In the case of the pure GtL or HVO oil phases, storage on water for two weeks resulted in the absence of water enrichment (Figure 6, center). In contrast, the oil phases with an admixture of 50% HVO as well as pure HVO showed an increased acid number after two weeks of storage on water and on water with microbes (Figure 12 center).

The trace element analysis of the oil phases confirmed the contamination of the pure GtL and HVO oil phases with additional phosphorus sources (0.4 and 0.2 mg/kg, respectively). In contrast, the nitrogen analysis showed that the fossil fuel oil HEL3 contains far more nitrogen (54.1 mg/kg) than the pure GtL or HVO (14.4 mg/kg and 15.8 mg/kg, respectively) (Figure 7).

The composition of GtL and HVO was identified by GC x GC/MS and accurately quantified by GC x GC/FID (Table 4 top and Figure 13, respectively). It was found that GtL and HVO consist exclusively of n-/iso alkanes. The alkanes have a chain length range restricted to C11- C20 in the case of HVO, and a broader chain length range of C8-C25 in the case of GtL.

### 3.5 Microbial activity below future blends of paraffinic fuels and biodiesel

#### CO_2_-Monitoring (1:5 approaches)

To analyze the possible effect of the future combination of alternative paraffinic fuels (GtL or HVO) and biodiesel (RME or UCOME) in heat supply on microbial activity or susceptibility, a 14-day storage culture series was run with 20% RME plus 50% GtL or 50% HVO as oil phase. The comparison was made to cultures with fossil heating oil HEL2 or fossil heating oil with 20% RME. Under fossil heating oil, microbial activity measured by the sum of CO_2_ accumulation in 300 ml headspace and 250 ml oil phase after 14 days was 15 mg. Under fossil heating oil with the addition of 20% RME, CO_2_ accumulation of 54 mg was possible, corresponding to a 269% increase in microbial activity. The further addition of 50% HVO did not significantly affect the microbial activity (48 mg CO_2_). In contrast, the addition of 50% GtL allowed only 34 mg CO_2_ microbial activity after 14 days, corresponding to a 127% increase in microbial activity compared to fossil heating oil (Figure 15A).

**Figure 15.**
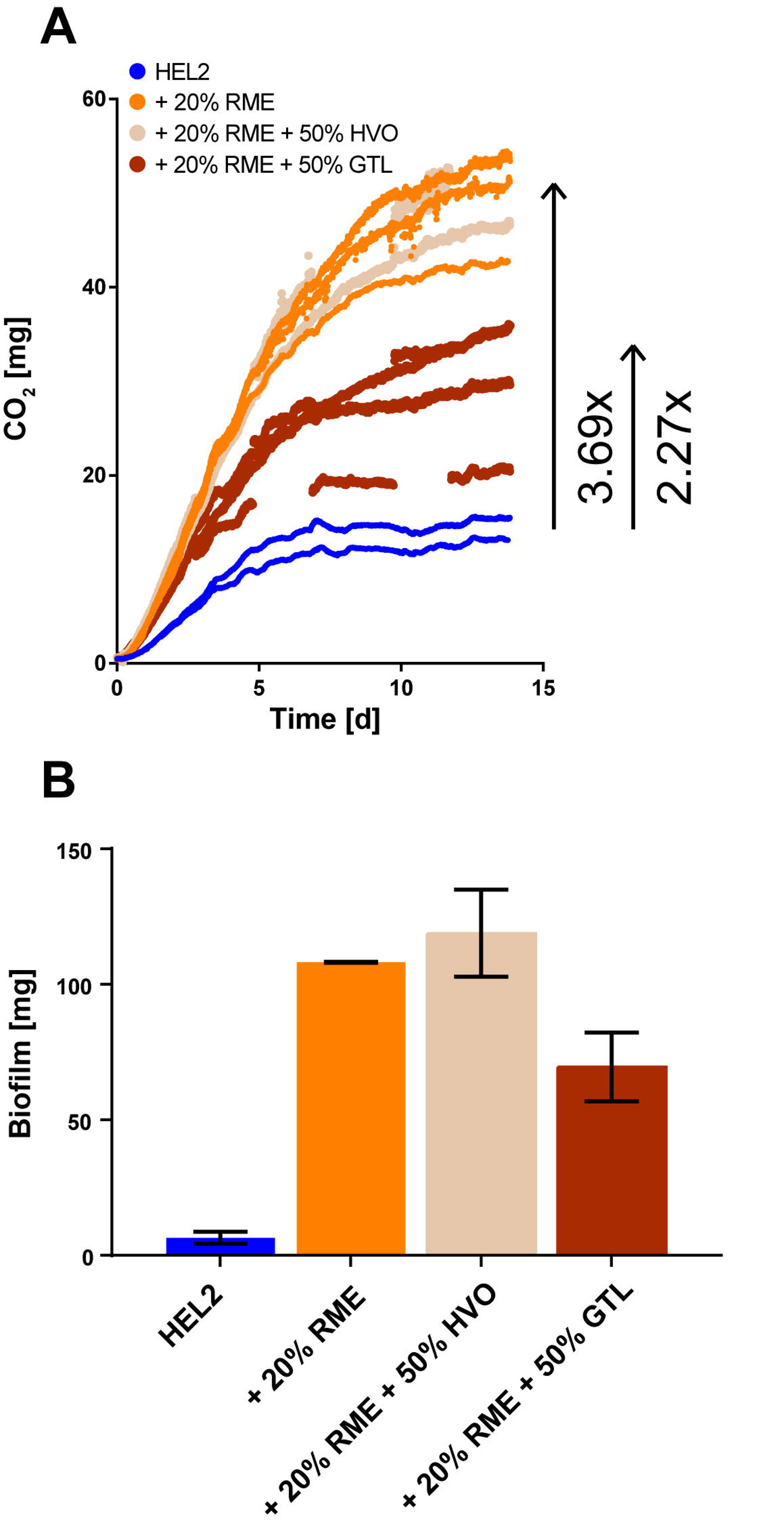
Storage cultures of RME/HVO and RME/GtL blends. Microbial activity in a 14-day storage culture (1:5) with fossil heating oil HEL2, fossil heating oil with 20% admixture of RME, with 20% admixture of RME plus 50% admixture of HVO, and of 20% admixture of RME plus 50% admixture of GtL is shown. Microbial activity is measured by the sum of CO_2_ accumulation in oil phase and headspace. Plotted is the discontinuous CO_2_ measurement of up to four biological replicates. Plotted are up to four biological replicates (A). The mass of biofilm formed in 14-day storage cultures (5:1) with fossil heating oil HEL2, fossil heating oil with 20% admixture of RME, with 20% admixture of RME plus 50% admixture of HVO, and of 20% admixture of RME plus 50% admixture of GtL is shown. Mean values and standard deviation of two technical measurements of pooled samples from each of four biological replicates (B) are plotted.

#### Biofilm formation (5:1 approaches)

In addition to monitoring microbial activity, as measured by the accumulation of CO_2_, the possibility of biofilm formation was quantitatively compared depending on the oil phase used. For four pooled biological replicates, 6 mg of biofilm was harvested after 2 weeks for the fossil heating oil HEL2. For the addition of 20% RME or 20% RME plus 50% HVO, a 17- fold and 18-fold increase in measured biofilm to 108 and 119 mg, respectively, was recorded.

In contrast, for the addition of 20% RME plus 50% GtL, only an 11-fold increase to 70 mg biofilm could be documented (Figure 15B).

## 4. Discussion

During the time window of both types of storage cultures (1:5 and 5:1 approaches), the onset of microbial contamination was simulated. This includes selection processes of the microbial mixture used and initial biofilm formation at the interface between the water and oil phases. The production of microbial biosurfactants, emulsion formation, and the increasing spread of microbes into the oil phase in the course of advanced contamination by an adapted microbial mixture was not mapped in this work.

In addition, it has to be noted that a pre-existing water phase was used in this work, since the presence of a free water phase is crucial for the progression of microbial contamination (Figure 1) [17, 59].

### 4.1 Fossil heating oils of six different German refineries define the baseline of microbial activity

A 14-day storage culture series (1:5) comparing potential microbial activity with fossil heating oils from six different German refineries revealed considerable variation between HEL4 products with 12 mg or HEL2 with 14 mg accumulated CO_2_ and HEL1 or HEL3 products with 26 mg CO_2_ each (Figure 2A). Comparatively reduced surface tensions and pH values indicated acidic and surface-active extracts from the oil phase of HEL1 and HEL3 (Figure 2B and C), which could be considered as bioavailable nutrient sources. A corresponding HPLC-IR analysis for their detection was not successful. Compensation of pH and surface tension in the presence of microbes (Figure 2B and C, +MO) indicated that these potential nutrients are utilized on the one hand and were not available in unlimited quantities on the other. This influence on surface tension also directly affects the water content of the oil phase. For the untreated heating oils, HEL1 and HEL3 with higher contents of extractable acidic and surface-active substances, a naturally higher water content was also determined. When the oil phases are incubated on water without microbes, the substances from the oil phase reduce the surface tension of the water phase, and the water content of the oil phases increases (Supplementary Figure 1). However, the applied microbes degrade the surface-active substances and an increase in the water content in the oil phase does not occur or the water content decreases again (incubation on water with microbes or incubation on water plus incubation on water with microbes). In this scenario of beginning contamination in a heating oil storage, the microbes thus stabilize the surface tension and limit further mass transfer instead of improving access to the potential nutrient source by producing their own surfactants.

A gas chromatography-tandem mass spectrometry (GC-MS/MS) analysis of water-soluble components of the heating oils indicated that in storage cultures the water phases among the tested heating oils were predominantly enriched with mono-aromatic compounds for which acute toxic effects on microorganisms have been documented. Deviating from this, the heating oils HEL3, HEL5, and HEL6 resulted in higher proportions of di-aromatic compounds, which are characterized less by acute toxicity but increasingly by carcinogenic properties [116]. A higher prevalence of di-aromatic compounds may correlate with a reduced occurrence of benzene, again suggesting a possible reduced acute toxicity (see chapter 3.1). However, these are only initial indications based on literature findings; for a reliable statement on the toxicity of the individual fuel oils in comparison to each other, microbial toxicity tests would have to be carried out.

However, to demonstrate the resulting differences in the toxicity of the water extracts, these water extracts were used as water phase in additional storage culture series (1:5) under the corresponding fossil heating oils (Figure 3). The demonstration of a 37% reduction in microbial activity when using the water extracts from heating oil HEL2 indicated that the comparatively higher toxicity of the water-soluble components was one reason for the lower microbial activity under heating oil HEL2 compared to other fossil heating oils.

The fossil heating oils are not only a source of toxic substances against which the microbes have to stand, they are also the source of all relevant nutrients. In this respect, the nitrogen analysis revealed crucial differences between the fossil heating oils. In the heating oils HEL1, HEL3, and HEL5, which allow the highest microbial activities (Figure 2A), the three highest concentrations of bound nitrogen were detected at the same time (Figure 7). A supplementary storage culture series, in which 2.9 g/L of additional ammonium chloride was added to the water phase under heating oil HEL3, showed that microbial activity below fossil heating oils as the only nutrient source is nitrogen-limited (Supplementary Figure 3). However, the heating oil with the highest nitrogen content did not automatically allow the greatest microbial activity (Figure 2A and 7). The nitrogen content does not exclusively determine the possible microbial activity. The further GC x GC/MS analysis (Figure 8) also showed that the nitrogen sources contained in the fossil heating oil HEL3 were challenging polyaromatic heteroatomic compounds such as carbazoles.

As examples of fossil heating oils that allow increased or low microbial activity, the products HEL3 and HEL2 were examined in detail by GC x GC/MS and GC x GC/FID without treatment and after storage on microbes (5:1). In the case of the heating oil HEL2 (Figure 8), the addition of a paraffin oil consisting of n-/iso- and cyclo-alkanes led, on the one hand, to a reduced water content of the oil phase (Figure 6 left) and, on the other hand, to an underrepresentation of the aromatics (Table 3A and C). The additional quantification of the degradation performance on both oil phases as a result of two weeks of storage (5:1) showed that the lower proportion of aromatics permanently reduced the attractiveness of the heating oil HEL2 as a carbon source measured concerning the total degradation performance on both heating oils. This is because all three possible carbon sources (n-/iso alkanes, cyclo alkanes, and aromatics) were utilized to the same extent in both heating oils by the microbial mixture used here (Table 3B and D). The efficient degradation of (poly)aromatic (heteroatomic) hydrocarbons has so far been described primarily for bacterial species (e.g. *Pseudomonas sp.*, *Acinetobacter sp.*, *Micrococcus sp.*), and bacterial species are highly adaptable to these demanding nutrient sources [117–120]. Also in the storage cultures of this study, a dominance of bacterial species was found as a sequencing result (Figure 5 left) for the water phases among all fossil heating oils, indicating the predominant utilization of aromatics as an important carbon and possible nitrogen source by bacteria.

### 4.2 Toxic degradation products of OME blends prevent microbial activity

The influence of oxymethylene ethers (OME) on microbial activity was investigated by a storage culture series (1:5) with an admixture of 2-10% OME to fossil heating oil HEL2. The admixture of OME provided complete suppression (85%) of microbial activity in the storage culture (Figure 9A). The parallel lowering of pH and surface tension and the subsequent HPLC analysis of control samples without microbes (1:5) (Figure 9B-D) showed that the admixed OME decomposes in contact with the free water phase to its starting products formaldehyde and methanol. Subsequent oxidation of formaldehyde also provided small amounts of formic acid sufficient to lower the pH of the unbuffered water phase. By repeating the storage culture series with buffered water phase (phosphate buffer) and stable pH (data not shown) and by compensating the pH when 2% OME was added to the oil phase by microbes present (< 10 mM formic acid, Figure 9C), it was shown that the pH effect is not crucial for inhibition of microbial activity. The integrity of the OME used for blending was subsequently confirmed by GC analysis (Supplementary Figure 2). The long-term storage cultures with a 10% OME admixture to fossil heating oil HEL2 (Figure 10) and regular measurement of extractable gDNA as a measure of the biomass present also showed that the antimicrobial effect of formaldehyde and methanol had persisted for at least one year and that no rapid adaptation of the broad microbial mixture took place. In the fuel sector, the application of oxymethylene ethers as a fuel alternative with a maximum admixture of 15 %(v/v) is aimed at, and the application as a fuel additive or performance improver for cleaner combustion is discussed [93, 95–99]. It seems obvious to exploit or commercialize the additional antimicrobial or preservative effect of OMEs or their decomposition products in contact with infiltrated water, as identified here. Given the ongoing registration with the European Chemicals Agency (Cas No. 30846-29-8 or 66455-31-0) and the ongoing preparation of a standard for the power and fuel sector (DIN/TS 51699), it is important to declare the antimicrobial effect as a side effect and to avoid classification as a biocide under the BPR (Biocidal products regulation). The classification as a side effect is supported by the fact that the antimicrobial effect only comes into play from an addition of 1-2 % (v/v) to the fuel. The typical use of biocides in the fuel sector is < 100 ppm. The existing registration of the released formaldehyde as a biocide (CAS No. 50-00-0) and the possible violation of occupational exposure limits for formaldehyde of 0.37 mg/m^3^ when released into the gas phase have to be taken into account for regulatory classification and application, though.

### 4.3 Intensification of the microbial activity below rapeseed oil methyl ester and used cooking oil methyl ester blends

For the biodiesel variants (RME and UCOME) used in this section, GC x GC/MS and GC x GC/FID analysis was performed to identify compositional differences as a reason for divergent effects on microbial activity. Small shifts to shorter chain lengths in UCOME compared to RME indicated that the used cooking oil was not pure rapeseed oil (Table 4). An admixture of palm oil with an average lower chain length range (palm oil: C16-C18, rapeseed oil: C18-C22), which has taken larger shares in the production of biogenic power and fuel in Germany in recent years [121], is likely. The observation of additional ethyl esters in UCOME (Table 4 and Figure 13 below) showed that methanol with traces of ethanol was used in the methylation. No effect on microbial activity was expected or observed for the documented deviations of UCOME from RME.

A two-and-a-half-week storage culture series (1:5) provided a twofold accumulation of microbially produced CO_2_ for blending the market standard 20% biodiesel (RME and UCOME) to fossil heating oil HEL2. Oil phases from pure biodiesel also allowed only two and a half times the accumulation compared to fossil heating oil. In addition, a bending of the CO_2_ curve could be observed for all approaches after 14 days at the latest. Thus, although a clear increase in microbial activity or catabolic activity of the developing biomass could be observed by the addition of biodiesel, as was to be expected based on existing literature [12, 14, 16, 18, 33; 57-59], there was also a technical limitation of the measurement technique used (Figure 11A).

While the storage of half of the produced CO_2_ in the oil phase has already been pointed out as a technical limitation in the previous work and has already been considered here, the oxygen limitation [37] and the occurrence of leaks at the culture vessel at higher CO_2_ concentrations come into question as additional limitations. Regardless of a possible technical limitation, it is striking that in the first days of microbial contamination in heating oil storage, the extent of microbial activity is independent of the amount of biodiesel added. In both cases, the attractive nutrients of the biodiesel are initially available in unlimited quantities. Only in the course of a contamination process lasting several months or several years, differences in the progress of contamination due to shortage of these nutrients can be expected with the addition of 20% biodiesel.

As in the case of fossil heating oils, the addition of biodiesel to fossil heating oil HEL2 (1:5) resulted in a strong reduction in surface tension and pH in the water phase (Figure 11B and C). As the cause of the reduction in surface tension, methanol, the starting material for methyl ester production, was detected by HPLC analysis with a surface tension of 23 mN/m (Figure 11D). In contrast to the observations for the fossil heating oils, the presence of microbes and their degradation activity did not lead to a re-stabilization of surface tension and pH, which indicated a significantly higher availability of the additional simple carbon sources (Figure 11 B and C, +MO Week 1 and 2). In contrast, for an oil phase of pure UCOME, the microbes were able to stabilize the surface tension. HPLC analysis of the water phase provided higher amounts of glycerol instead of methanol, a by-product of methylation, as one reason for this. Glycerol has a surface tension of 60 mN/m.

In the oil phases with higher proportions of biodiesel and thus higher content of polar simple carbon sources, a continuous increase in water content was already observed without further treatment (HEL2: 21 mg/kg, 20% UCOME: 96 mg/kg, 100% UCOME: 264 mg/kg, 20% RME: 179 mg/kg, 100% RME: 534 mg/kg) (Figure 6 right). By storing the oil phases on water (0.1% NaCl), a further increase in the water content takes place, depending on the addition of biodiesel and in parallel to the reduction of the surface tension in the water phase, independent of the presence of microbes. This increase in water content is less pronounced in the case of UCOME, which corresponds to the stabilized surface tension. In the case of the addition of 20% RME, 100% RME, and 100% UCOME, the measured water contents exceed the theoretical water absorption capacity of the oil phase (depending on the biodiesel content: in the case of 20% biodiesel 200 mg/kg, in the case of 100% biodiesel 1500 mg/kg), the formation of emulsions is likely, the better access to the nutrients of the oil phase and the better dispersion even in the oil phase is possible for the microbes [19, 11–13, 16–17, 59].

As expected, the trace element and nitrogen analysis of the oil phases showed contamination with boron, phosphorus, and nitrogen for the biodiesel additions, which are typical components of plant fertilizers. This resulted in an optimization of the C:N:P ratio in the case of both biodiesel types and thus an optimized decomposition performance of the carbon sources. However, the simple inorganic phosphorus and nitrogen sources are accessible to all microbial species, in contrast to the observations for the fossil heating oils. The additional documentation of silicon in the case of UCOME indicates the addition of silicone oil. This is a common antifoam agent used in the oil industry, which was the second reason for the stabilized surface tension when an oil phase of pure UCOME was used in the storage cultures (Figure 7 below).

The addition of biodiesel ultimately resulted in a variety of additional simple carbon sources (free fatty acids, fatty acid methyl esters, glycerol, methanol, and others) as well as phosphorus and nitrogen sources accessible to all microbial species, such that sequencing of the complete water phases from the storage cultures (1:5) for the addition of biodiesel documented an immediate expansion of microbial diversity to include the yeasts and molds (Figure 5 right). This has immediate implications for the complexity and robustness of biofilms formed, as well as the metabolic repertoire of the contaminant running off in terms of degradation capacity and microbially assisted corrosion [123–126].

### 4.4 Limited microbial activity below Gas-to-Liquids and hydrogenated vegetable oils

The influence of the alternative paraffinic fuels was investigated using hydrogenated vegetable oil (HVO) and a gas-to-liquid (GtL), a representative of the Fischer-Tropsch products (XtL). In a first storage culture series, the comparison was made with the fossil heating oil HEL3, which was one of the fossil heating oils with the highest microbial activity in the previous sections (24 mg CO_2_ in this storage). Blending the market standard 50% HVO resulted in a 28% reduction in potential CO_2_ accumulation in comparison, while blending 50% GtL and pure HVO and GtL resulted in a 45% reduction to 13-15 mg CO2 (Figure 14A). This corresponds to the minimum microbial activity previously observed for fossil heating oils (see Figure 2A).

In parallel, under oil phases with the paraffinic alternatives, a continuous stabilization of the surface tension of the water phases (Figure 14B) and, associated with this, a reduction of the water content in untreated oil phases as well as an absence of water accumulation in the storage cultures (Figure 6 middle) could be observed. This corresponds to lower availability of surface active and potentially simple nutrients from the oil phase. There is also a lower likelihood of emulsions and microbial dispersal into the oil phase. While stabilization of the pH in the water phase was observed for the admixture of GtL (Figure 14C), the admixture of HVO provided slightly increased acid numbers in the oil phase due to storage on water (Figure 12 middle). The hydrolysis-induced release of fatty acids in the case of HVO is certainly due to the plant feedstock, but is by no means comparable to the extent in the case of biodiesel blending (Figure 12 right) and did not lead to increased microbial activity for the time being.

The blending of GtL and HVO resulted in an increase in phosphorus content according to trace element analysis and, although not more nitrogen overall (14-15 mg/kg for pure GtL and HVO, respectively), additional simple inorganic nitrogen sources. The fossil fuel oil HEL3 used for blending already contained large amounts (54 mg/kg) of polyaromatic nitrogen sources, as discussed in previous sections. Thus, the admixture of HVO and GtL now also allowed yeast and mold to achieve an optimized C:N:P ratio (Figure 7). Sequencing showed that the 50% admixture of HVO and GtL allowed increased microbial diversity by stabilizing the proportions of yeasts and molds in the presence of bacterial dominance (*Burkholderia sp.*) (Figure 5 middle). In contrast, a renewed restriction of microbial diversity was observed for oil phases of pure GtL and HVO. The crucial difference was the restriction to n-/iso alkanes as the only carbon source and the omission of aromatics. Nitrogen and phosphorus continued to be readily available to all microorganisms. Thus, in addition to an optimized C:N:P ratio, the supply of species-dependent simple or diverse carbon sources is crucial for the expression of microbial diversity.

The 50% admixture of GtL to fossil heating oil HEL3 resulted in a stronger decrease in microbial activity than the 50% admixture of HVO (Figure 14A). Besides the hydrolysis-induced release of free fatty acids as simple carbon sources in the case of HVO, the GC x GC/MS or GC x GC/FID analysis of the oil phases provided further reasons (Table 4 top and Figure 13, respectively). The GtL used possessed a higher proportion of n-/iso alkanes of shorter chain length. Short-chain alkanes react non-specifically with the lipid membrane of microorganisms and have toxic effects [127–129]. To confirm the toxic effect of GtL, an additional storage culture series was run in which the additional admixture of 50% HVO or 50% GtL to fossil heating oil HEL2 with 20%RME was investigated (Figure 15A). The admixture of 20%RME resulted in an increase in microbial activity from 15 mg to 54 mg CO_2_. This result was unaffected by the addition of 50% HVO (48 mg CO_2_). Only the admixture of 50% GtL resulted in microbial activity of only 34 mg CO_2_ after two weeks. In this culture series in the classic 1:5 approach, however, the subjective impression also arose that the biofilm formation between the water and oil phases was not affected by the toxic effect, but on the contrary, was even stronger with the admixture of 50% HVO or 50% GtL. Since the formation of biofilm poses a crucial threat to the functionality of the burner systems, a third culture series was performed in a 5:1 format to harvest and weigh the biofilms afterward (Figure 15B). Quantification of the biofilms confirmed the CO_2_ monitoring results (microbial activity). Biofilm formation was unchanged by blending 50% HVO to 20% RME (108 mg and 119 mg, respectively). In contrast, biofilm formation was reduced to 70 mg by blending 50% GtL to 20% RME.

## 5. Conclusion

The addition of oxymethylene ethers revealed an antimicrobial side effect that could be activated by water, the comparison of fossil heating oils from different refineries showed a considerable range of microbial activity, the addition of biodiesel confirmed the acceleration of microbial contamination, and the addition of paraffinic fuels, although also partly of biogenic origin, proved to be microbially resistant alternative heating oils. This work shows that fossil heating oils and their CO_2_-reduced alternatives, whether biogenic or synthetic, impose multiple parameters on the onset of microbial contamination that promote, restrict, complement, or limit the microbial contamination process. Species-specific usable nitrogen sources of the oil phases promote microbial activity, while toxic extracts, such as formaldehyde or monoaromatics (BTEX), limit microbial activity. An optimized C:N:P ratio enhances microbial diversity, but is limited by demanding carbon sources. The use of advanced chromatography, microbial sequencing, and element (nitrogen and phosphorus) analysis can determine in detail the basis of an accelerated or retarded contamination process. Surface tension and pH or acid number and water content at the boundary between infiltrated water and oil phase, as well as CO_2_ accumulation in the storage tank headspace, can be used as indicators of the onset of microbial contaminations. With this analytical workbench, one can determine the susceptibility of heating oil to microbial contaminations and can use this information for prevention, for example by additive addition or fuel blending.

## Acknowledgments

The authors are grateful for the practical support provided by the Institute of Technical and Macromolecular Chemistry (ITMC) of RWTH Aachen University in the person of Mrs. Celine Jung for the GC analysis of the oxymethylene ether solution used.

## Authorś Contributions

## Funding

This work was supported by the German Federation of Industrial Research Associations “Otto von Guericke” e.V. (AiF) [IGF Nr. 20840 N, 2019]. The laboratory of Lars M. Blank was also partially funded by the Deutsche Forschungsgemeinschaft (DFG, German Research Foundation) under Germany’s Excellence Strategy within the Clusters of Excellence TMFB 236 and FSC 2186 ‘The Fuel Science Center’. The funding sources were not involved in the study design, data acquisition, analysis, interpretation, and the decision to submit the manuscript.

---

**Supplementary Figure 1. Influence of microbes on the water content of fossil heating oils during storage cultures.** The water content of HEL3 after storage for two weeks in 5:1 approaches on 0.1% NaCl plus microbes (A) and on 0.1% NaCl (water) (B) is shown. In addition, the water content of the untreated heating oil and the water content after storage for two weeks initially on 0.1% NaCl (water) followed by storage for two weeks on 0.1% NaCl plus microbes is shown (B followed by A). Plotted are individual measurements from up to 4 biological or technical replicates and the corresponding mean value.

**Supplementary Figure 2. GC analysis of the OME solution used.** Shown is that the mass of all OME components found in the sample is 94.9% of the weighed sample mass. The GC method does not take into account the OME component OME-6 contained in the sample, which, however, could be observed as a peak and explains the missing mass fraction.

**Supplementary Figure 3. Relevance of nitrogen for microbial activity in storage cultures of fossil heating oils.** Microbial activity in a 14-day storage culture of fossil heating oil HEL3 with a water phase of 0.1% NaCl and with a water phase of 0.1% NaCl plus 2.9 g/L NH_4_Cl is shown. Microbial activity is measured by the sum of CO_2_ accumulation in oil phase and headspace. Plotted is the discontinuous CO_2_ measurement of up to three biological replicates.

## Notes

### Competing Interest Statement

The authors have declared no competing interest.

