## Supplementary figures and images for "Evaluating microbial contaminations of alternative heating oils"

### Supplementary Figure 1

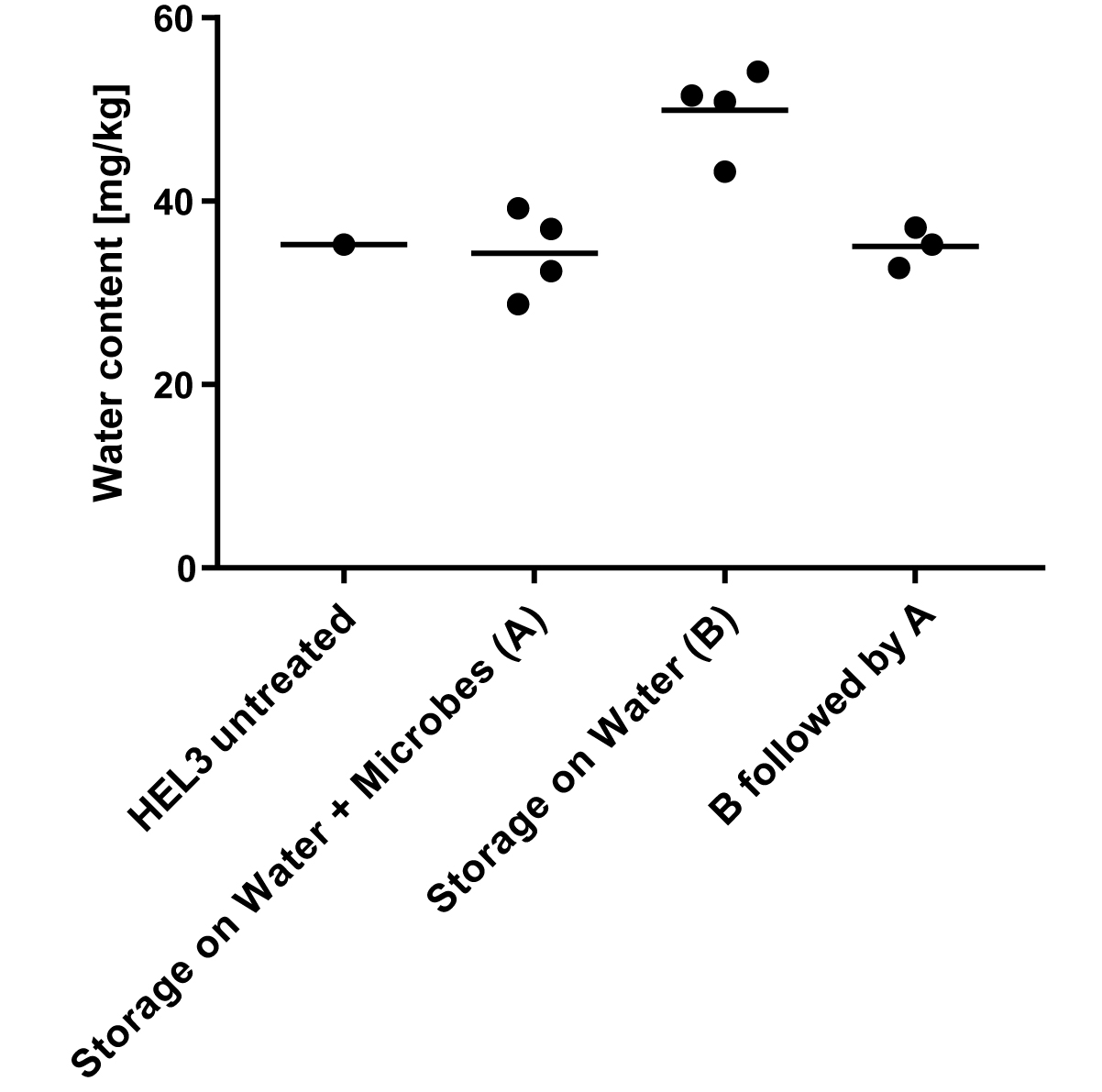

### Supplementary Figure 2

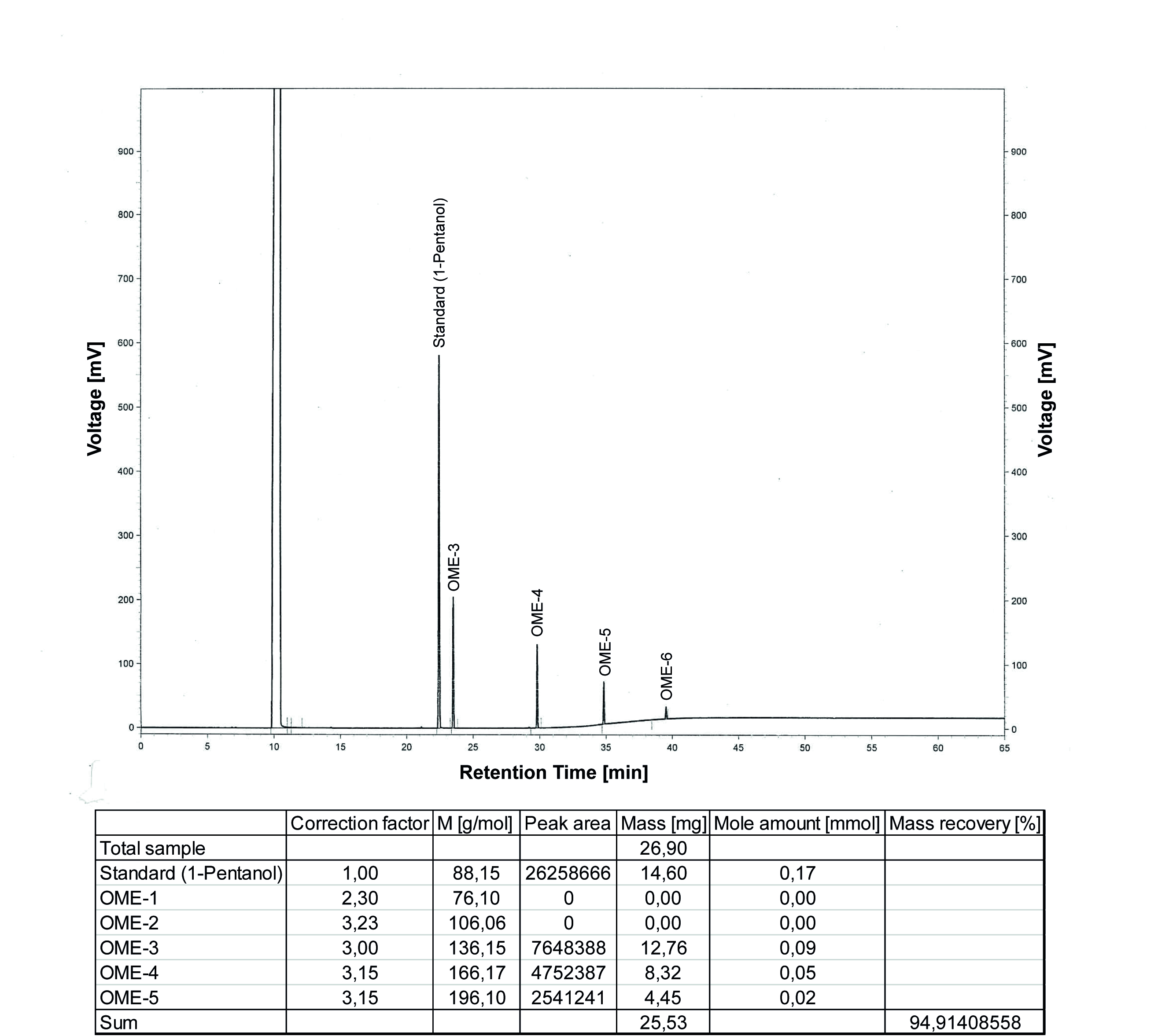

### Supplementary Figure 3

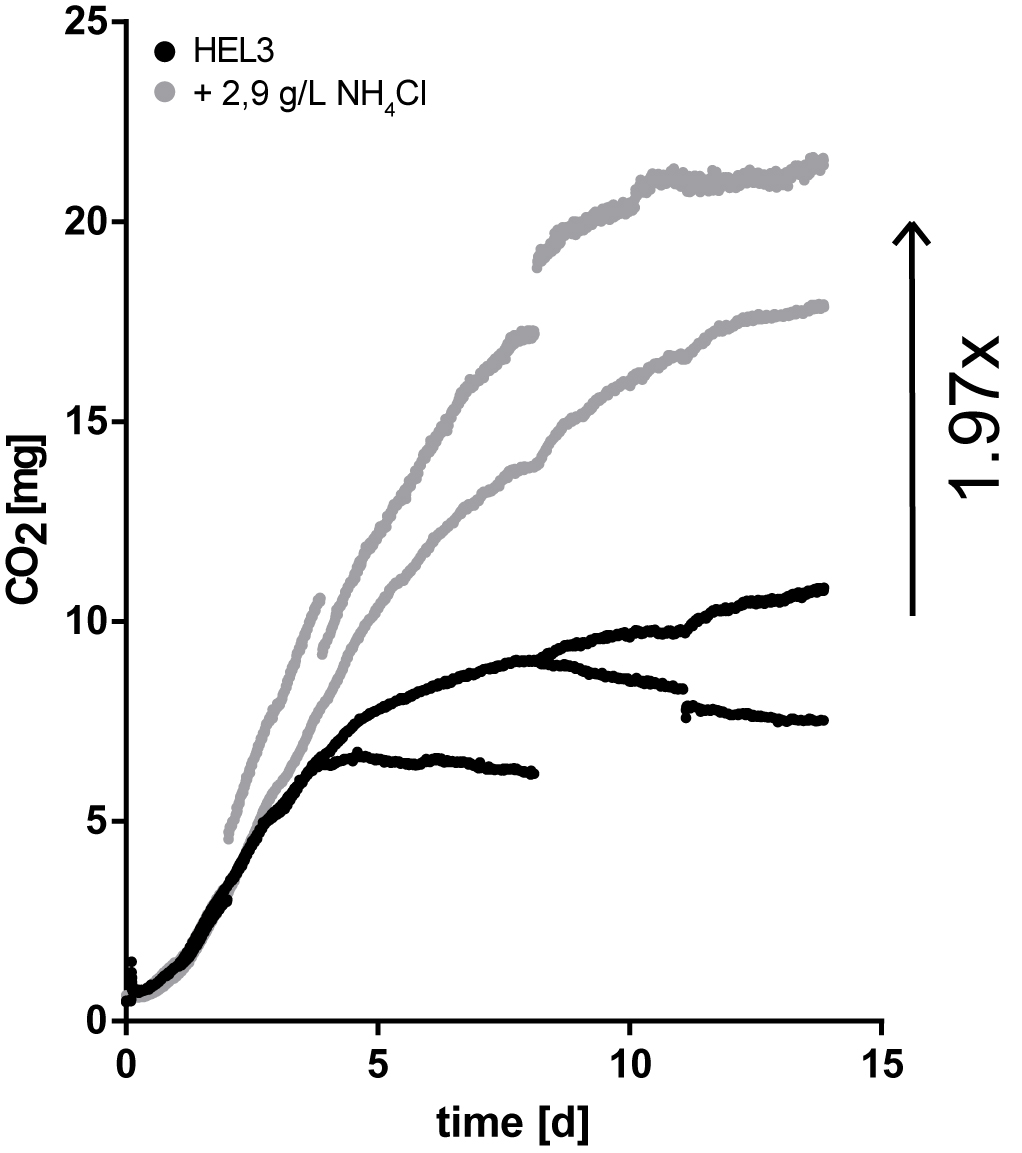
